# Neuron type-specific mRNA translation programs provide a gateway for memory consolidation

**DOI:** 10.1101/2025.01.13.632784

**Authors:** Mauricio M. Oliveira, Olivia Mosto, Robert Carney, Wendy J. Liu, Emmanuel Makinde, Catherine Leckie, Carson Schultz, Emily H. Lu, Karen S.A. Ruiz, Thomas J. Carew, Eric Klann

**Author notes:** Co-second authors (Olivia Mosto, Robert Carney, Wendy J. Liu).

## Abstract

Long-term memory consolidation is a dynamic process that requires a heterogeneous ensemble of neurons, each with a highly specialized molecular signature. Considerable effort has been devoted to identifying molecular changes that accompany the process of consolidation, but mostly hours or days after training, when memory consolidation is already complete. Studies have shown that protein synthesis is elevated during the early stages of consolidation, but how this increase impacts neuronal function remains unclear. We hypothesize that mRNAs translated during the early stages of consolidation could provide information on how diverse neurons involved in memory formation restructure their molecular signatures to support memory. Here, we generate a landscape of the translatome of three neuron types in the dorsal hippocampus during the first hour of contextual memory consolidation. Our results reveal that translation programs associated with consolidation are different among neurons, fueling the reconfiguration of specific biological processes. We further demonstrate the patterned translation of mRNAs in different neuron types during consolidation is explained by features hard-coded in the mRNA sequence, suggesting ubiquitous mechanisms controlling activity-induced neuronal translation. Altogether, our work uncovers previously unknown mechanisms controlling activity-induced translation in neurons and provides a large, readily available resource for scientists interested in the role of memory formation in health and disease.

## Introduction

The birth of a new memory involves the synchronous engagement of a heterogeneous population of neurons, promoting molecular changes that strengthen the connectivity of a neuronal ensemble in the brain^1^. These neurons can be excitatory or inhibitory, depending on the specialized molecular signatures and electrophysiological properties they carry^2–4^. The molecular state of a neuron enables it to quickly respond to converging afferent stimuli and average these signals into one final output. Different molecular states can drive varied methods of integrating stimuli to produce a wide range of responses. These profiles are not fixed, as they may be fine-tuned or even drastically changed in response to salient activity. However, there is a severe lack of understanding regarding the molecular dynamics that occur on short timescales in different neurons *in vivo*, especially in the context of memory formation. Most studies interrogate the molecular profile of neurons at long time points (e.g. days to weeks) after memory acquisition, when memory formation has already completed, but do not examine how this process of consolidation starts and develops. Furthermore, these studies generally rely on measurements of the transcriptome at the single-cell level, which is unlikely to capture true functional differences, given that single-cell sequencing generates sparse data enriched in highly-expressed transcripts, and transcript levels do not automatically correspond to protein abundance^5,6^. On the other hand, studying the dynamic changes in the proteome is challenging due to low technical sensitivity, especially when studying small neuronal subpopulations. At the intersection between transcriptome and proteome, the translatome, or the full set of translating mRNAs, represents a promising alternative approach to dissect functional changes occurring in neurons during consolidation.

The translatome is especially attractive given that long-term memory consolidation is characterized by rapid recruitment and transient enhancement of translation^7–11^, whose disruption is capable of modifying memory formation^12,13^. This reinforces translational mechanisms as potential therapeutic targets for memory-related disorders, such as post-traumatic stress disorder and Alzheimer’s disease. The translatome at early stages of memory consolidation has been examined previously^14^, but the lack of cell type specificity of these studies precludes the determination of differential mRNA programs triggered in discrete neuron populations involved in long-term memory consolidation. Furthermore, without cell type specificity, glial cell contamination serves as a significant confound even at the level of whole neuronal populations. On the other hand, when the translatome was studied in a neuron type-specific manner^11^, a weaker threat memory paradigm was utilized, resulting in smaller molecular changes. Given the critical role of *de novo* translation during memory consolidation, there is an imminent need to identify the neuron type-specific molecular changes that underlie stable memory formation.

Here, we leveraged translating ribosome affinity purification (TRAP)^15^ combined with RNA-sequencing to examine threat conditioning-induced alterations in the translatome of three different neuron types of the dorsal hippocampus (dHPC) known to be important for long-term memory consolidation: excitatory neurons expressing *Camk2a*^+^, and inhibitory neurons expressing *Pvalb^+^* or *Sst*^+^. We find that these neurons rapidly recruit specialized mRNAsto ribosomes (henceforth referred to as ribosome-associated mRNAs and abbreviated as RA-mRNAs). We further demonstrate a fundamental role for the translation initiation modulator GADD34 in translatome dynamics and long-term memory, revealing its role as an upstream regulator of conditioning-induced modifications to the translatome. Finally, we find that rapid modifications of the translatome during consolidation are a function of cis-elements embedded in the mRNA sequence, particularly AU-rich elements (AREs), and disrupting a candidate ARE-targeting RNA binding protein leads to robust memory deficits. Altogether, this work provides the scientific community with a new resource and extensive hypothesis-generating database that uses cellular and systems-level analyses aiming to unravel the molecular basis of long-term memory consolidation.

## Results

### Contextual threat conditioning is a robust method to study consolidation-associated neuronal *de novo* protein synthesis

We used contextual threat conditioning to gain an understanding of neuron plasticity events linked to memory consolidation. This behavioral paradigm is profoundly reliant on the dorsal hippocampus (dHPC)^16^, requires *de novo* protein synthesis^17–19^, and induces detectable molecular changes in neurons. In our experiments, mice that underwent the threat conditioning paradigm display increased rates of freezing (i.e. defensive) behavior as they are shocked, as opposed to their control counterparts (“Box”), that are presented to the new context but are not shocked (Ext Fig 1a, left). The next day, shocked mice displayed significantly higher freezing than control mice when exposed to the threatening context, confirming formation of an aversiveassociative memory (Ext Fig 1a, right panel). To exclude the possibility that any molecular changes observed in future experiments were simply induced by the foot shock, we designed an experiment where one group of mice underwent the traditional threat conditioning paradigm, and the other was subjected to a latent inhibition paradigm, where inconsequential pre-exposure to the context leads to weak associative learning post-exposure to shock^20^ (Ext Fig 1b). After exposure to the context, mice were injected retro-orbitally (R.O.) with azidohomoalanine (AHA) to label short timescale *de novo* protein synthesis induced by conditioning^21^. We found that the traditional threat conditioning paradigm induced a robust increase in neuronal *de novo* protein synthesis, but not when preceded by latent inhibition (Ext Fig 1c-d). However, all conditions induced an increase in the number of c-Fos^+^ (Ext Fig 1e, g) and phosphorylated ribosomal protein S6^+^ (S6P^+^) neurons in the dHPC (Ext Fig 1f, h), suggesting that, as expected, neurons are activated across all groups, but that *de novo* protein synthesis elevation is uncoupled from the expression of c-Fos^+^/S6P^+^. Overall, these results allowed us to draw two main conclusions: (1) changes at the translational level should be studied at large populational levels rather than isolated c-Fos^+^/S6P^+^ neurons; and (2) increases in *de novo* protein synthesis are not linked to foot shock administration, but rather to associative contextual learning.

**Figure 1.**
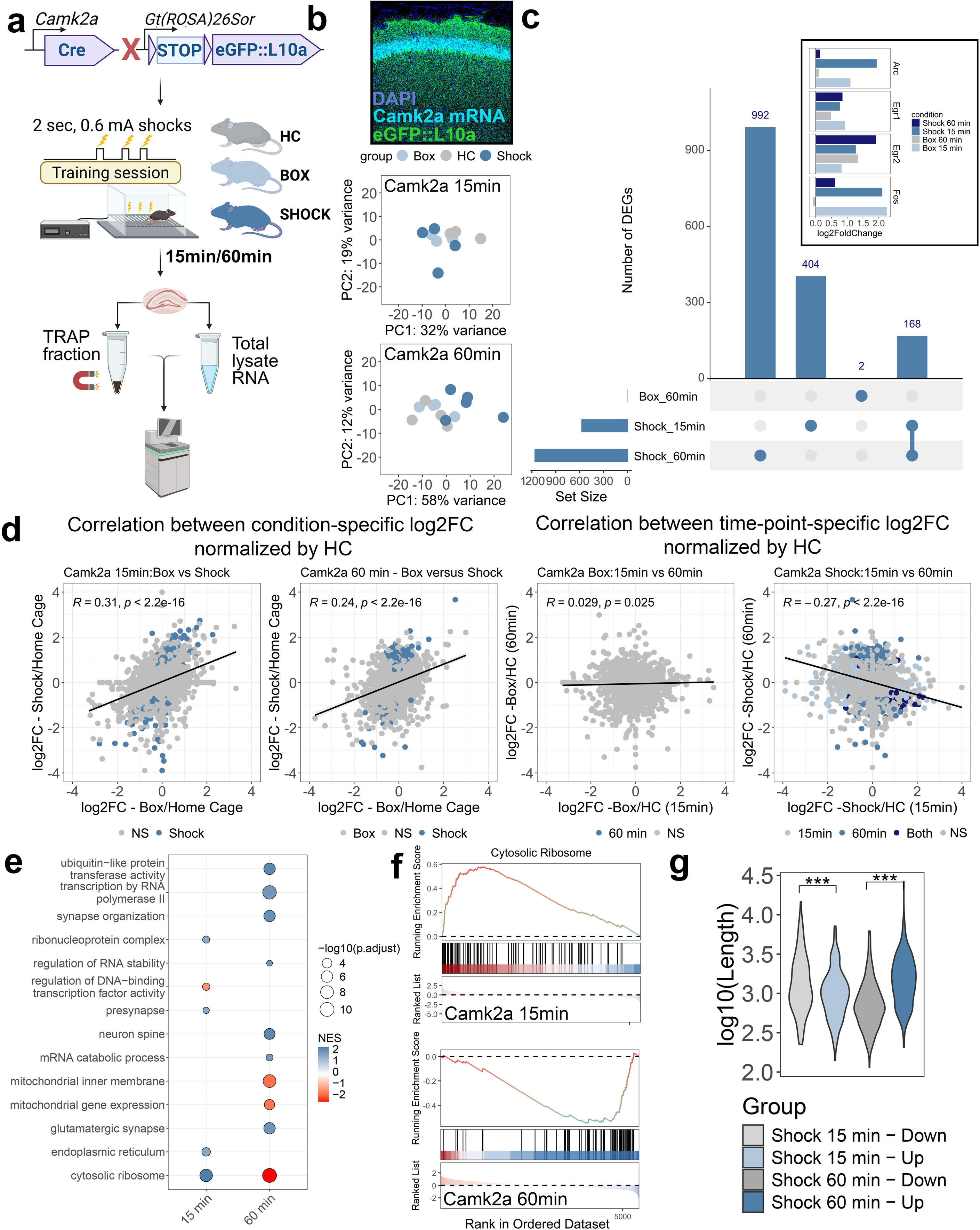
Examination of conditioning-induced short timescale changes to the translatome of excitatory neurons reveal a connection between RA-mRNAs CDS length and ribosomal occupancy. (a) Experimental design and workflow. (b) Upper panel. Co-staining of anti-GFP immunohistochemistry (green) and RNAscope for *Camk2a* mRNA (white). Blue is DAPI. Middle and lower panels: PCA of TRAP-sequencing samples of 15min and 60min datasets. *Home cage* = grey. *Box =* Light blue. *Shock* = dark blue. (c) Upset plot registering the intersection of DE RA-mRNAs found in different conditions across datasets. Numbers above bars represent total number of RA-mRNAs per intersection. Bar plot in lower left represents the total number of DE RA-mRNAs per group. Connected dots in bottom identify the intersections. Inset = Bar plot representing the differential expression of four IEGs (*Arc, Egr1, Egr2* and *Fos*) across conditions. (d) Correlation plots between conditions in each time point (two plots on the left) or between time points in each condition (two plots on the right). NS = non-significant. Light blue = significant at 15min time point. Blue = significant at 60min time point. Dark blue = significant at both time points. At the top left of each plot, *R* is the Pearson’s correlation coefficient, *p* value is the correlation significance. (e) Gene ontology analysis of DE RA-mRNAs identified in *shock* condition of each time point. Blue = Upregulated terms. Red = Downregulated terms. Size is -log10 adjusted *p* value. (f) Gene set rank plot for GO “*Cytosolic ribosome*”. Each vertical line represents the position of a RA-mRNA belonging to the GO in ranked gene list. Upper plot represents the 15min time point, lower plot represents the 60min time point. (g) Violin plots comparing the coding sequence lengths of up- or downregulated RA-mRNAs in each time point. Y axis is log-transformed length of CDS. *** = *p <* 0.001. Student’s *t*-test.

### Examination of conditioning-induced short timescale changes to the translatome of excitatory neurons confirms a connection between RA-mRNA CDS length and ribosomal occupancy

To gain insight into molecular changes occurring in neurons during the initial stages of long-term memory consolidation, we leveraged translation ribosome affinity profiling (TRAP). This technique enables efficient recovery of cell type-specific translating ribosomes bound to their client mRNAs through overexpression of ribosomal protein L10a (RPL10a) conjugated to eGFP, which is targeted with anti-eGFP antibodies^15^. Subsequently, RA-mRNAs are purified using oligo(d)T beads and sequenced through conventional RNA-sequencing techniques.

Increased mRNA translation in excitatory neurons following conditioning is associated with threat memory formation. This phenomenon was previously described in both the dHPC^11^ and the amygdala^12,13^, suggesting that increased *de novo* protein synthesis is a ubiquitous feature of memory consolidation. Therefore, we investigated threat conditioning-induced changes to the translatome of excitatory neurons residing in the dHPC by conducting contextual threat conditioning in *Camk2a*.Cre-TRAP^fl^ mice (Fig 1a; Ext Fig 2a), which express RPL10a::eGFP only in excitatory neurons (Fig 1b, upper panel). To gain insight into the dynamics of the translatome during the first hour of consolidation, we collected samples 15 or 60 min after conditioning (Fig 1a). Home cage mice (“HC”, i.e. never introduced to the conditioning paradigm) and mice that went inside the box but were not shocked (“Box only”), were included as controls.

**Figure 2.**
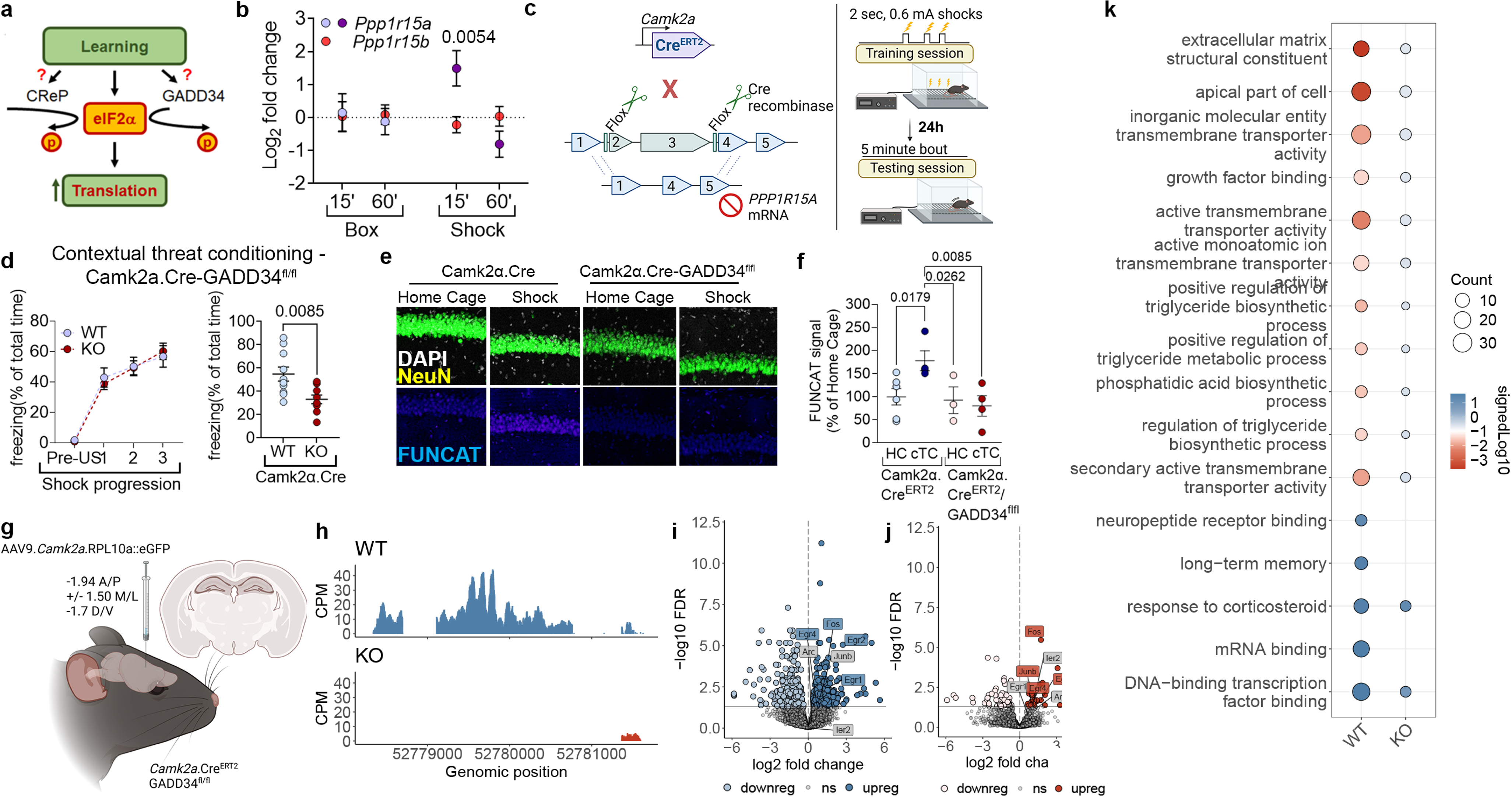
GADD34 controls fast-paced changes to the pool of RA-mRNAs induced by threat conditioning. (a) Rationale for eIF2α dephosphorylation-mediated increase in *de novo* translation. (b) Log_2_FC of *Ppp1r15a* (mRNA encoding GADD34) and *Ppp1r15b* (CReP) in the translatome of *Camk2a^+^* neurons of mice that underwent contextual threat conditioning against home cage baseline*. \*\** is *p =* 0.0054; Number of mice per group in the dataset used is listed in *Methods* section. Mean ± SEM. (c) Schematics of contextual threat conditioning paradigm to test the relevance of GADD34 expression in excitatory neurons in long-term memory formation. (d) Contextual threat conditioning of *Camk2a.Cre ^ERT^*^2^*/GADD34^flfl^.* Left panel = training session. Right panel = testing session. *N =* 9-10 mice/group; Mean ± SEM; *F* = 2.871. (e) Representative images of dynamic labeling of *de novo* protein synthesis in the dHPC of *Camk2a.Cre ^ERT^*^2^*/GADD34^flfl^.* Top panels: white = DAPI; yellow = NeuN. Bottom panels: blue = FUNCAT. (f) Quantification of (E). Two-Way ANOVA with Dunnet’s post-hoc correction. Mean ± SEM; *F =* 4.079. (g) Experimental plan for TRAP-seq of excitatory neurons from the dHPC of *Camk2a.Cre^ERT^*^2^*/GADD34^flfl^* mice. *N* = 3-5 mice/group. Mean ± SEM. (h) Track plot of read coverage found in the gene encoding *Ppp1r15a*, demonstrating absence of GADD34 translation in knockout mice. (i) Volcano plot illustrating the DEGs found in WT. Light blue = downregulated. Dark blue = upregulated. Grey = not significant. (j) Volcano plot illustrating the DEGs found in KO. Light red = downregulated. Dark red = upregulated. Grey = not significant. (k) Gene ontology of differentially expressed RA-mRNAs found in WT and KO mice. Size = gene counts per gene set. Color = signed -log10 *p* adjusted.

Our TRAP-sequencing analyses identified ∼5900 RA-mRNAs enriched in ribosomes of excitatory neurons, following a strict filtering process (see *Methods*). Principal component analyses (PCA) revealed a very mild change in the translatome 15 min after conditioning (Fig 1b, middle panel), but these changes became clearer at the 60 min timepoint (Fig 1c, lower panel). Indeed, differential expression analyses revealed 9.67% and 19.65% of significantly modified RA-mRNAs in the *shock* group, when compared against the *home cage* group (Suppl Table 1). The same analyses comparing *box only* and *home cage* found 0% and 0.03% of differentially expressed RA-mRNAs, indicating that the negative valence associated with contextual memory induces more robust changes to the translatome. We compared our findings at the 60 min timepoint with a previous RiboTag experiment using similar conditions^11^. We found a 4-fold increase in differentially expressed RA-mRNAs amount (Ext Fig 2b). We attribute this discrepancy to conditioning intensity (3x 0.6mA foot shocks in our settings versus 1x 0.35mA foot shock paired with a tone).

We next interrogated whether the changes to the translatome were consistent across the two time points investigated. Thus, we examined the degree of overlap between the differentially expressed RA-mRNAs found in different time points. Overall, we found little overlap between the two time points in the *shock* condition, with only 168 RA-mRNAs significantly altered in both datasets (Fig 1c). Nonetheless, it is noteworthy that regardless of condition, there was marked increase in the expression of immediate early genes (IEGs) when compared to baseline (Fig 1c, inset), though not statistically significant in the *box* conditions. This suggests that the hippocampus has been activated by exposure to a new context, regardless of foot shock administration, consistent with our findings in Ext Fig. 1.

Presentation to a new context leads to dHPC activation, but we could not detect significant genes in our *box* datasets. Thus, we were interested in determining whether the two tested conditions (*box* and *shock*) led to similar overall trends in differential expression of RA-mRNAs. A correlation analysis identified a significant positive correlation between the two conditions at both 15 and 60 min (*R* = 0.31 and 0.24 for 15 and 60min, respectively), suggesting that presentation to a new context can lead to similar but diminished patterns of translatomic changes seen after threat conditioning (Fig 1d, two left panels). Interestingly, similar analysis comparing timepoints in each specific condition yielded different results, with almost no correlation in the *box* condition, and a negative correlation in the *shock* condition (Fig 1e). Overall, our results demonstrate that the RA-mRNAs in *Camk2a*^+^ cells dynamically change throughout the first hour of consolidation, with the immediate response (15 min) differing considerably from the late-stage response (60 min).

To examine the biological properties overrepresented in the differentially expressed RA-mRNAs, we filtered significantly altered genes (*p adj* < 0.05) and performed gene ontology analyses (Fig 1e; Ext. Fig 2e-f to examine their ranked position; Suppl Table 2). Our results demonstrate that ribosomal protein (RP)-encoding mRNAs are significantly enriched at 15 min but depleted at 60 min (Fig 1f). The decrease in RP-encoding mRNAs at 60 min was not observed in the previous RiboTag dataset^11^, where a weak training paradigm was utilized (Ext Fig 2c-d). Further, we found that mRNAs related to neuronal plasticity are enriched only at 60 min, together with mRNAs encoding transcription- and proteostasis-related proteins. Finally, we found that ribosomes were depleted of RA-mRNAs encoding mitochondrial metabolism-related mRNAs.

The rapid translation of RP-encoding mRNAs led us to hypothesize that the time-dependent effects in the translatome were influenced by the coding sequence (CDS) length. Short CDS mRNAs, usually encoding housekeeping genes^22^, are more rapidly translated than long ones, such as those encoding synaptic proteins^23^. Consistent with this notion, we found that mRNAs upregulated at 15 min have significantly shorter CDS compared to downregulated mRNAs, a pattern that is reversed at 60 min (Fig 1g). This confirms a role for CDS length as a regulator of activity-induced neuronal translation and provides an explanation for the inverse correlation found between the two time points (Fig 1d).

### GADD34 controls rapid changes to the pool of RA-mRNAs induced by threat conditioning

The results described above motivated us to investigate a molecular mechanism that may control rapid shifts in RA-mRNAs induced by conditioning. We recently identified the protein phosphatase 1 co-factor GADD34 as responsible for promoting activity-induced translation of numerous plasticity-related mRNAs *in vitro*^24^. GADD34 (*Ppp1r15a*) is an inducible regulator of translation that, along with its constitutive counterpart CReP (*Ppp1r15b*)^25^, shares the task of promoting dephosphorylation of the eukaryotic initiation factor 2 on its alpha subunit (eIF2α). We reasoned that one of these proteins could be responsible for mediating conditioning-induced changes in RA-mRNAs (Fig 2a). We found in our translatome dataset that GADD34 translation is upregulated in excitatory neurons 15 min after conditioning (Fig 2b), which was not the case for CReP. These findings are consistent with our previous results *in vitro* and directed us to target GADD34 as a critical modulator of translation during memory consolidation.

The phosphorylation status of eIF2α has been demonstrated to have a bidirectional effect on memory. Its accumulation leads to memory impairment, whereas its ablation results in long-term memory enhancement^11,26,27^. Although a loss-of-function mutation in CReP leads to memory impairment, this is due to chronic elevation of eIF2α phosphorylation in the brain rather than specifically preventing the conditioning-induced elevation of *de novo* protein synthesis^28^, which is consistent with the known role of CReP as a constitutively active cofactor. On the other hand, the inducible nature of GADD34 led us to hypothesize that, similar to eIF2α phosphorylation status, GADD34 modulation would have a bidirectional effect on long-term memory formation. We first generated mice harboring a deletion of GADD34 in excitatory neurons and tested their memory using contextual threat conditioning (Fig 2c). We found that the mice presented no learning problems during the training session but displayed significantly less freezing behavior during the test session, indicating an impairment in long-term memory (Fig 2d). On the other hand, virally overexpressing GADD34 in excitatory neurons in the dHPC (Ext Fig 3a) led to long-term memory enhancement on the test day (Ext Fig 3b-c). These results demonstrate that, similar to eIF2α phosphorylation, expression of GADD34 in excitatory neurons has a bidirectional effect on long-term memory formation.

**Figure 3.**
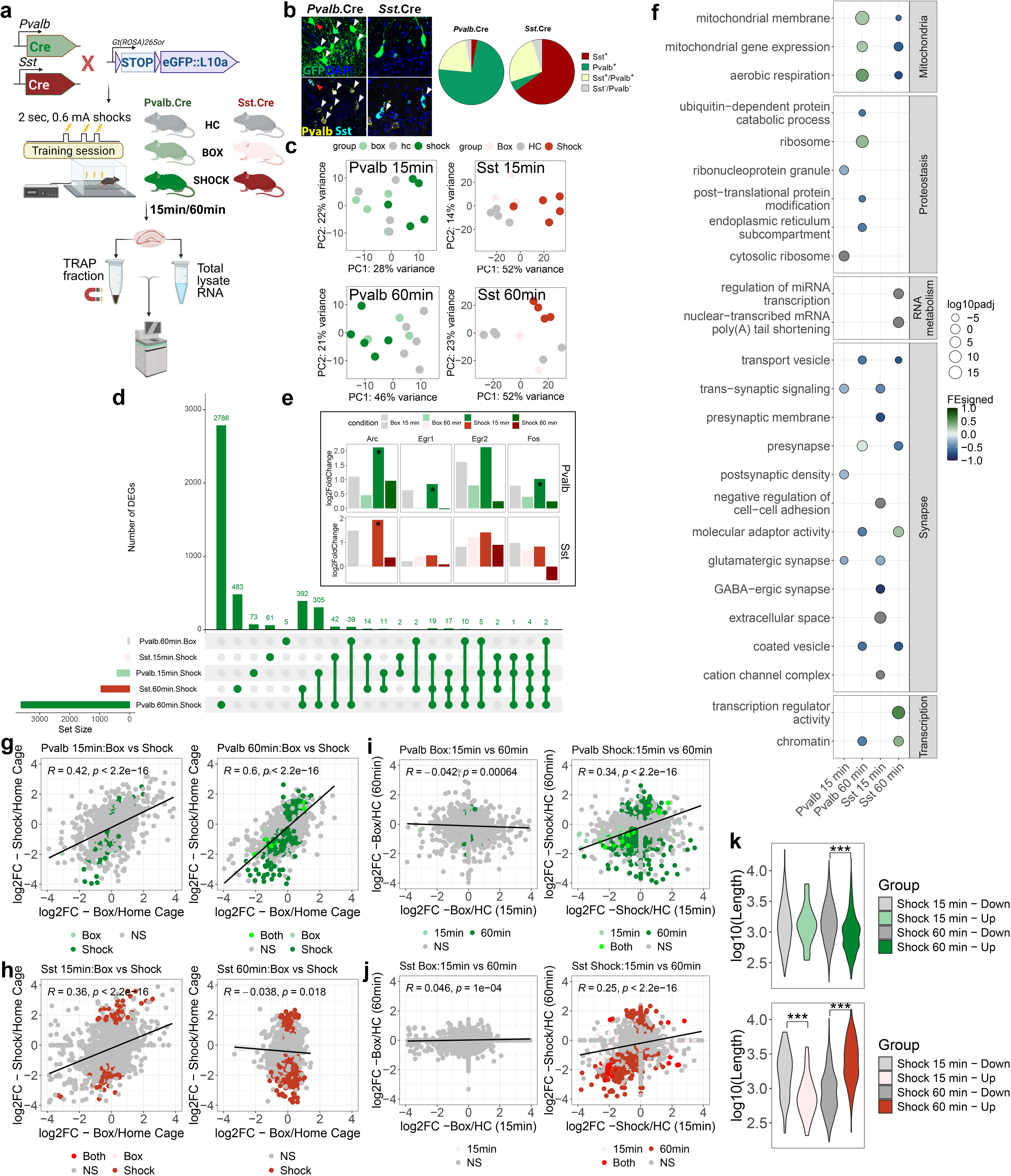
Conditioning-induced changes to the translatome of *Pvalb^+^* and *Sst^+^* interneurons reveal cell-specific patterns. (a) Experimental design and workflow. (b) Multiplex RNAscope exploring the overlap between *Sst^+^* and *Pvalb^+^* neuron populations in the dHPC. Green = GFP immunostaining; Yellow = *Pvalb*; Cyan = *Sst*. Blue = DAPI. White arrows exemplify expected cell type with no overlapping RNAscope signal from the other cell marker. Red arrow demonstrates an example of a cell carrying both cell markers. Right panel: pie plot with quantification of neuronal markers among GFP^+^ cells. Dark red = *Sst^+^;* Green = *Pvalb^+^*; Yellow = *Sst^+^/Pvalb^+^*; Grey = GFP^+^, no markers. N=2 mice per genotype. (c) PCA of TRAP-sequencing samples of 15min and 60min datasets. *Home cage* = grey. *Box =* Light colored. *Shock* = dark colored. Left panels = *Pvalb^+^* datasets; Right panels = *Sst^+^* datasets. (d) Upset plot registering the intersection of DE RA-mRNAs found in different conditions across datasets. Numbers above bars represent total number of RA-mRNAs per intersection. Bar plot in lower left represents the total number of differentially expressed RA-mRNAs per group. Connected dots in bottom identify the intersections. (e) Bar plot representing the differential expression of four IEGs (*Arc, Egr1, Egr2* and *Fos*) across conditions. Asterisks represent significantly changed conditions. (f) Gene ontology analysis of mRNA programs triggered by threat conditioning. Two columns on the left are *Pvalb* datasets (15 and 60min), and the two on the right are *Sst* datasets. Color is signed fold enrichment. Size is -log10 p adjusted values. (g-h) Correlation plots between conditions in each time point. NS = non-significant. Light colors = significant at 15min time point. Dark colors = significant at 60min time point. Bright colors = significant at both time points. At the top left of each plot, *R* is the Pearson’s correlation coefficient, *p* value is the correlation significance. (i-j) Correlation plots between time points in each condition. NS = non-significant. Light colors = significant at 15min time point. Dark colors = significant at 60min time point. Bright colors = significant at both time points. At the top left of each plot, *R* is the Pearson’s correlation coefficient, *p* value is the correlation significance. (k) Violin plots comparing the coding sequence lengths of up- or downregulated RA-mRNAs in each time point. Y axis is log-transformed length of CDS. *** = *p <* 0.001. Student’s *t*-test (intra time-point comparisons only).

We then tested whether conditioning-induced increases in *de novo* protein synthesis would be prevented by the knockout of GADD34. We trained mice in the contextual threat conditioning paradigm, and performed R.O. administration of AHA, as above. We found that deletion of GADD34 blocked conditioning-induced increases in *de novo* protein synthesis in the dHPC (Fig 2e), suggesting that disrupted *de novo* protein synthesis underlies the long-term memory failure in these mice. We further noted that GADD34 deletion did not induce significant effects to baseline neuronal translation, consistent with the idea that its role is bound to neuronal activity, as was previously reported *in vitro*^24^. These results also suggest that there would be a broad impairment of the RA-mRNA program recruited in response to conditioning in the GADD34 knockout mice. To examine this, we injected AAV9.Camk2a.RPL10a::eGFP in the dHPC of Camk2α.Cre^ERT2^/GADD34^fl/fl^ mice, overexpressing the TRAP system in excitatory neurons (Fig 2g; Suppl Table 3). As genotype controls, we included Camk2α.Cre^ERT2^ mice. We then trained mice in the contextual threat conditioning and collected dHPC samples for TRAP-sequencing 60 min after training (Fig 1). As a control, we included *home cage* mice. Read coverage assessment confirmed the absence of GADD34 mRNA in the knockout dataset (Fig 2h). Differential expression analysis demonstrated that although control mice displayed a similar response to what was observed in Fig 1, GADD34 knockout mice had a reduced response, with approximately 1.2% of RA-mRNAs differentially expressed (Fig 2i-j). These findings indicate that there is a general failure in the recruitment of mRNA programs necessary for memory consolidation in the GADD34 knockout mice. Indeed, gene ontology analysis revealed no relevant enrichment of terms in the GADD34 knockout dataset (Fig 2k).

Finally, given their diametrically opposing effects to translation, we examined whether knocking out either phospho-eIF2α and GADD34 would lead to inverse effects on the translatome. Unexpectedly, genotypical comparison (i.e. wild type versus GADD34 knockout in our dataset; wild type versus phosho-dead eIF2α^11^) revealed that GADD34 knockout had an effect on RA-mRNAs that was not the inverse of that of eIF2α (Pearson’s *R =* -0.17; Ext Fig 3d). Gene ontology analysis indicated that 9 terms were inversed in the genotypes, most of which related to neuron development and neurogenesis, suggesting that tight control of eIF2-dependent initiation is required to balance expression of transcriptional programs and neuronal differentiation (Ext Fig 3e). Overall, our data provides strong evidence that the GADD34/eIF2α signaling axis orchestrates a conditioning-induced increase in *de novo* translation of mRNAs related to transcriptional and neuronal plasticity, which are required for proper long-term memory consolidation. Furthermore, it demonstrates that translatome studies can be used as a reliable resource for identifying key molecules involved in long-term memory consolidation.

### Conditioning-induced changes to the translatome of *Pvalb^+^* and *Sst^+^* interneurons reveal cell-specific patterns

The robustness of our findings in excitatory neurons led us to expand our dataset, probing two other neuron types of the dHPC—*Pvalb*^+^ and *Sst*^+^ interneurons—that exert local control over excitatory neurons, thereby critically regulating long-term memory formation. *Pvalb*^+^ cells are generally located in the granular layer, projecting to the soma of pyramidal cells; *Sst*^+^ interneurons, on the other hand, are located in the outer layer of the dHPC, *stratum oriens*, but project axons to the molecular layer, where excitatory neuron dendritic trees are present^29^.

We used the same experimental design as with excitatory neurons, by crossing *Pvalb* or *Sst.*Cre driver mouse lines with *RPL10a::eGFP*^fl^. These mice were then exposed to contextual threat conditioning and dHPC samples were collected 15 or 60 min after conditioning for TRAP-sequencing (Fig 3a; Ext Fig 4a). Because both neuronal populations were previously demonstrated to overlap^30^, we performed RNA scope coupled with anti-GFP immunohistochemistry and found that GFP expression was largely contained to the desired cell population in each mouse line (Fig 3b, upper panel).

**Figure 4.**
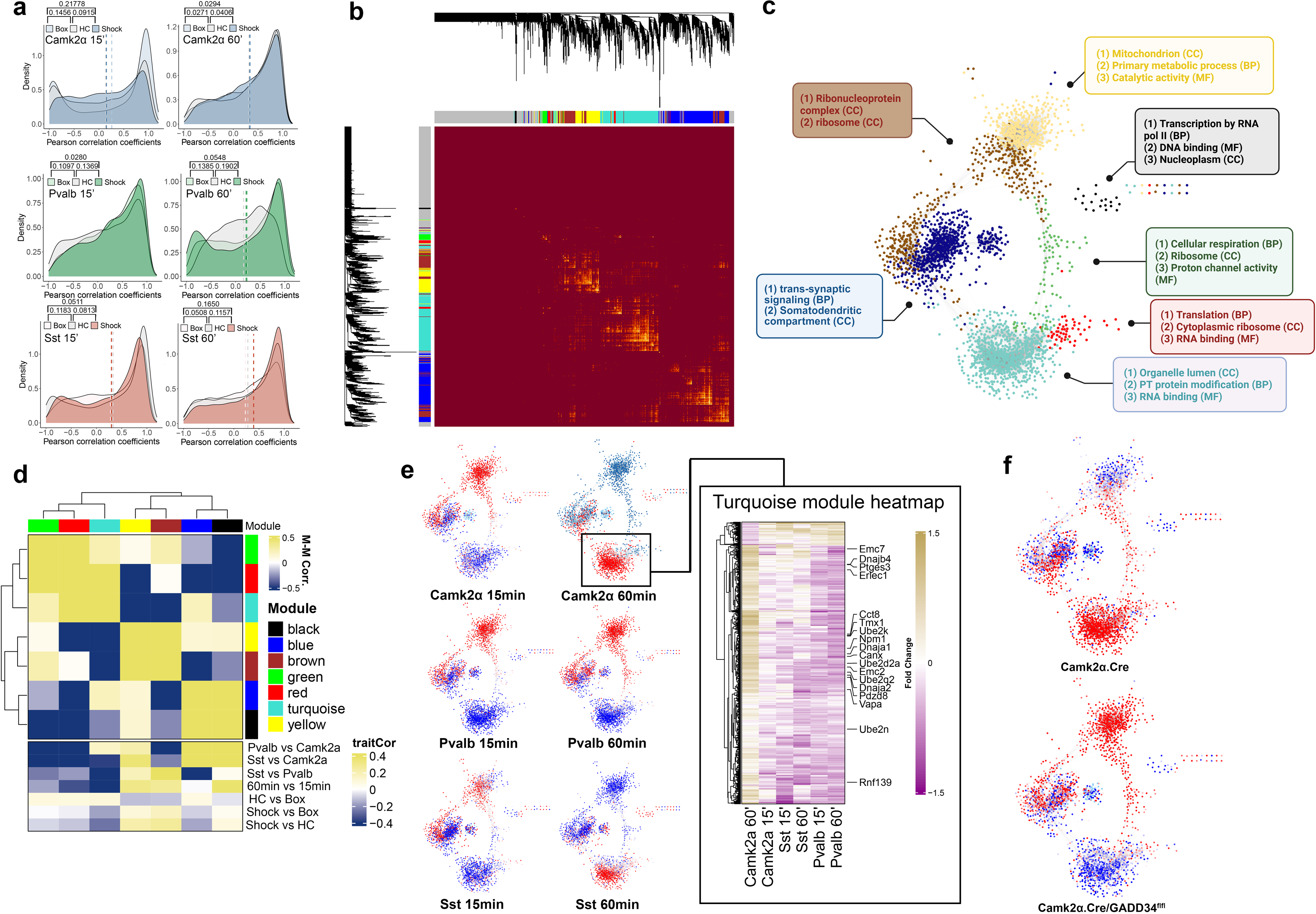
Global network analysis of conditioning-induced changes to neuronal translatome reveals the recruitment of specialized molecular programs by different neuron types. (a) Comparison of co-expression patterns from genes encoding proteins belonging to same protein complexes in *Camk2a*, *Pvalb* and *Sst* datasets (upper, middle and lower rows, respectively). X axis represents the distribution of correlation coefficients (Pearson’s *R*). Values on top left indicate the Wasserstein distance between Pearson’s *R* distribution along the x axis. Dashed vertical lines represent the arithmetic mean value of each distribution. (b) Pairwise correlation of expression patterns of RA-mRNAs across conditions. Bar graphs on the x and y axis represent the module assigned to each gene. Yellow regions represent high correlation values. Note that higher correlation values are generally contained to intra-module comparisons. (c) Representation of WGCNA network and the modules identified. Colors of modules are mapped to the network. Insets represent best overrepresented gene ontologies for each of the modules identified. CC = Cellular Component; BP = Biological Processes; MF = Molecular Function. e and module-trait correlation heatmap. Colors on the x and y axis of the main heatmap represent the modules. Color is module-module correlation. On the lower heatmap, color represents module-trait correlation. Each trait comparison is described to the right of each row, with the “*trait1 vs trait2”* consensus. High correlation values indicate better correlation with *trait1*, and negative correlation indicates better correlation with *trait2*. (e) Map of the conditioning-induced fold in the gene network obtained through WGCNA. Red = upregulated (25% upper IQR), blue = downregulated (75% lower IQR). Top left = *Camk2a^+^* neurons 15 minutes after conditioning. Top right = *Camk2a^+^* neurons 60 minutes after conditioning. Middle left = *Pvalb^+^* neurons 15 minutes after conditioning. Middle right = *Pvalb^+^* neurons 60 minutes after conditioning. Bottom left = *Sst^+^* neurons 15 minutes after conditioning. Bottom right = *Sst^+^* neurons 60 minutes after conditioning. Inset: Heatmap of fold changes of all mRNAs clustered in the blue module. Magenta = downregulated genes; Gold = upregulated genes. Selected mRNAs with symbols highlighted are encompassed in GOs “*Proteasomal protein catabolic process*”, “*Endoplasmic reticulum membrane”* and *“Unfolded protein binding*”. (f) Map of the log_2_FC values in the WGCNA network previously obtained. The WT network is similar to what was previously found, but the KO network is drastically different. Red = upregulated (25% upper IQR), blue = downregulated (75% lower IQR).

Initial assessment using PCA demonstrated a separation between the *shock*, *box* and *home cage* groups, at both 15 and 60 min (Fig 3c), suggesting significant alterations in the translatome induced by conditioning. After applying the same filtering process as in excitatory neurons, we identified ∼6500 RA-mRNAs enriched in *Pvalb*^+^ datasets, and *∼*4000 *in Sst^+^* cells (Suppl Table 4). In *Sst^+^* cells, we found similar fractions of DE RA-mRNAs as in excitatory neurons in the two timepoints: 2.78% at 15 min and 24.09% at 60 min. However, in *Pvalb^+^* cells, we identified a much larger fraction of differentially expressed RA-mRNAs: 6.42% at 15 min and 54.47% at 60 min. Again, *box* controls displayed minimal fractions of differentially expressed RA-mRNAs (0.03% and 0.2% at 15 min in *Pvalb*^+^ and *Sst^+^*, respectively; 0.9% and 0% at 60 min). Interestingly, intersection analysis revealed a shared backbone between the two interneuron sub-types in the conditioning-induced translatome response, with 392 shared DE RA-mRNAs at 60 min (Fig 3d). This, however, was not true at the 15 min time point (only 2 shared genes). Again, regardless of condition, we observed an increase in the expression of immediate early genes in both *box* and *shock* conditions, although it only reached significance at the 15 min timepoint of the *shock* condition (Fig 3e).

We next assessed the biological properties that were differentially modulated by conditioning in each neuron type. Gene ontology analysis identified mitochondrial metabolism as an early target in *Pvalb*^+^ interneurons (Fig 3f; Ext Fig 4b-e to examine their ranked position; Suppl Table 5), but that class of RA-mRNAs was downregulated in *Sst*^+^ cells. Interestingly, *Sst*^+^ cells showed an increase in factors involved in miRNA transcription, as well as poly(A) tail shortening factors, such as *Tnrc6a*, *Tnrc6b* and *Tnrc6c*, which encode proteins that act in synchrony with proteins from the Argonaute family to promote miRNA-dependent mRNA decay^31,32^. This suggests that *Sst*^+^ interneurons increase mRNA instability during early phases of consolidation, though the reasons for this remain unclear. Both *Pvalb^+^* and *Sst*^+^ neurons underwent changes at the synaptic level, most notably depletion of RA-mRNAs encoding proteins related to neurotransmission. These findings suggest that each class of interneurons have unique mRNA programs that are recruited to ribosomes after conditioning.

Given our findings that a similar translational response in excitatory neurons occurs in both *box* and *shock* groups, albeit with different magnitudes, we examined whether similar patterns could be found in interneurons. Our results suggest an overall conserved response in both interneurons, similar to what was found in excitatory neurons (Fig 3g-h), with the exception being *Sst*^+^ cells at 60 min, where no correlation was found between *box* and *shock* groups. When we compared the responses across time points, we found a significant positive correlation in the *shock* group (Fig 3i-j), which is an inverse of the pattern observed in excitatory neurons. Additionally, no correlation was observed within the *box* group. These findings suggest that interneurons recruit mRNA programs that are persistent throughout the first hour of consolidation, differing from the pattern of recruitment observed in excitatory neurons.

In excitatory neurons, we found a correlation between translation speed and CDS length. However, due to the differences in translation program dynamics between these neurons and their inhibitory counterparts, we hypothesize that CDS length would have a differential impact on translation speed in interneurons. In *Pvalb*^+^ interneurons, there appears to be no direct correlation between CDS length and timescale of ribosomal occupancy (Fig 3k, upper panel), but *Sst*^+^ neurons displayed similar patterns as excitatory neurons (Fig 3k, lower panel). This finding suggests that different neuron types of the dHPC employ varied mechanisms to control the translation of mRNAs during consolidation, which culminates in neuron type-specific conditioning-induced translatome programs.

### Gene regulatory network of conditioning-induced changes to neuronal translatome reveals the recruitment of specialized molecular programs by different neuron types

Given the challenging nature of identifying the mechanisms underpinning the distinct translational dynamics observed in these neuronal populations through individual examination, we opted to utilize an integrative approach. We determined whether the baseline translatome of the three neuron types would reflect well-known biological differences between the cells. PCA revealed three individual sample clusters representing individual neuron types (Ext Fig 5a). However, the interneuron clusters were more proximal than the excitatory cluster, suggesting a higher degree of shared identity in their translatome. We further found congruent enrichment of classical neuron markers (Ext Fig 5b). Excitatory neurons displayed enrichment of *Camk2a* and *Slc17a7*, whereas interneurons displayed enrichment of *Gad1* and *Gad2*, which encode enzymes of the GABA synthesis pathway. Furthermore, we found a significant enrichment of *Pvalb* in *Pvalb*^+^ datasets, and of *Sst* in *Sst^+^* datasets, indicating a clear separation of the two neuronal populations. Finally, we observed significant depletion of the glial markers *Aldh1l1, Gfap, Cnp* and *Cx3cr1*.

**Figure 5.**
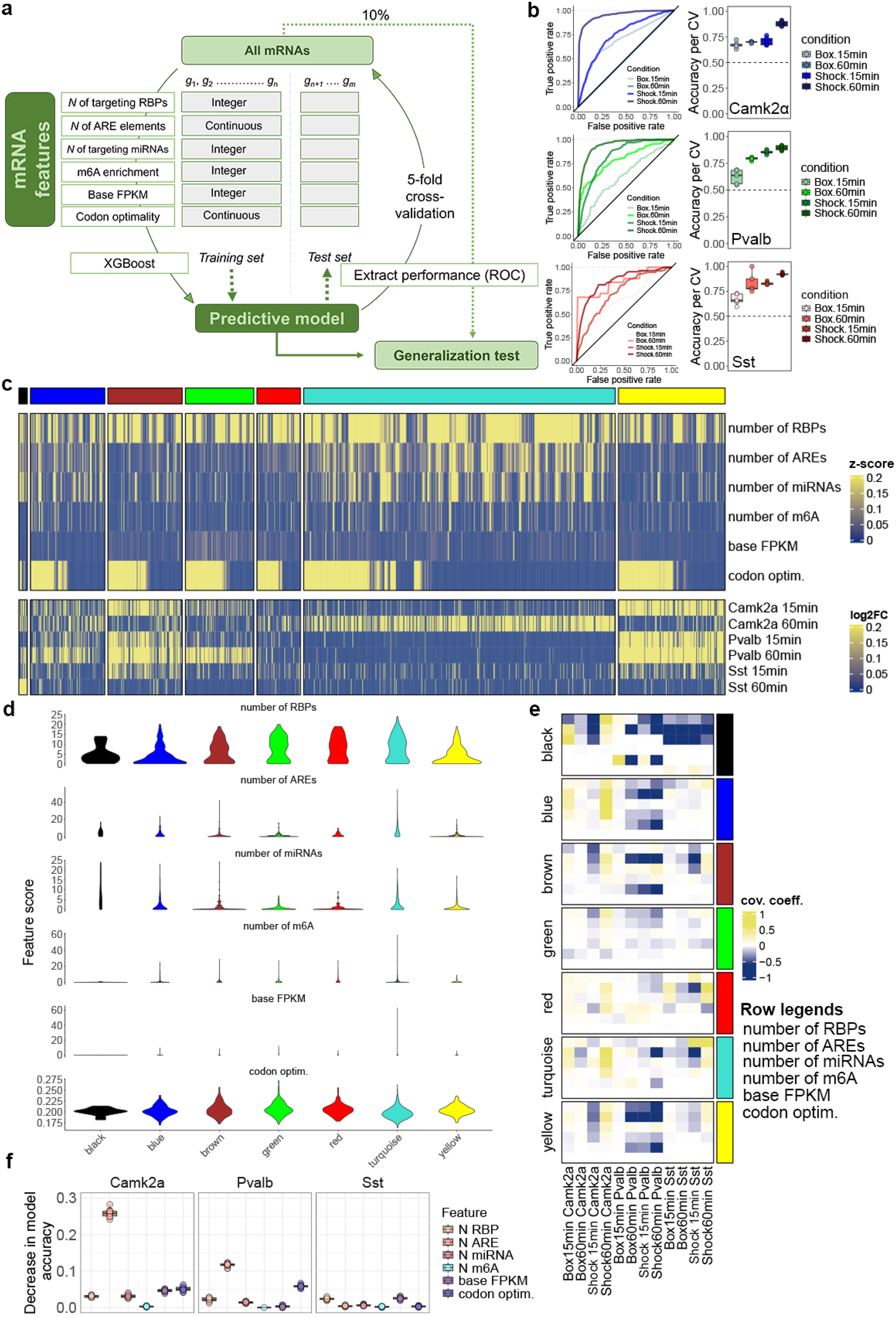
Cell-specific patterned changes in RA-mRNAs induced by conditioning are controlled by abundance of regulatory elements present in the 3’UTR of mRNAs. (a) Representative illustration of the design of the machine learning model. Adapted from ref^56^. (b) On the left: Receiver Operating Characteristics (ROC) curves on the left, categorized by condition tested. Diagonal line represents random classification performance (50% chance). On the right: Accuracy per cross validation performed (*k* = 5). Horizontal dashed line represents random classification performance. Blue shades graphs = *Camk2a*; Green shades graphs = *Pvalb*; Red shades graphs =*Sst*. (c) Heatmap depicting the distribution of mRNA features across GRN modules. Column annotation on the top represents each module built by WGCNA. Rows in the main heatmap represent normalized mRNA features. Lower heatmap depicts the fold changes found in all *shock* datasets, compared against *home cage*. Bottom annotation represents the module-trait correlation, only against time points. Yellow = high expression; Blue = low expression. (d) Violin plots depicting the distribution of features across modules. Feature score (y axis) represents the absolute values of each feature found in each module. Statistical comparisons can be found in supplementary table 9. (e) Co-variance matrix between conditions found in our dataset and feature presence. Row splits represent each module (depicted by the right-sided annotation). Columns represent each condition. In each row split, each row represents an individual feature (legend at the bottom right of the heatmap). (f) Permutation importance test (*k* = 5).

After our filtering process (see *Methods*), we determined that 15.5% of RA-mRNAs overlap among all neuron types, whereas 52.9% were enriched in only one neuron type (Ext Fig 5c). Differential expression analysis comparing each interneuron sub-type versus excitatory neurons revealed that interneurons display similarities in their translational programs when compared to excitatory neurons (Ext Fig 5d; Suppl Table 6). However, we were still able to determine that 27.0% of RA-mRNAs were differentially expressed among both interneurons. Gene ontology analysis found enrichment of terms known to be associated with each cell type, such as *“glutamatergic synapse*” for *Camk2a*^+^ cells and *“extracellular matrix*” for interneurons (Ext fig 5e; Suppl Table 7). We further found that excitatory neurons display enrichment of mRNAs associated with RNA biology when compared to inhibitory neurons, including RP-encoding mRNAs, rRNA pre-processing and ribosomal assembly. Overall, the direct comparison of the neuron-specific translatomes confirmed that we can identify conserved features classically associated with each neuron type, allowing for their direct comparison.

Proteins belonging to the same functional unit were previously shown to have correlated abundance in neurons^33^. We thus asked whether this pattern was also reflected in the pool of mRNAs associated with ribosomes. We found that RA-mRNAs encoding proteins of the same functional units have higher correlation of expression when compared to a random sampling from the original pool of RA-mRNAs in the *home cage* group (Ext. Fig 6). However, conditioning did not significantly change correlation of expression patterns between mRNAs encoding related proteins, suggesting that this does not explain conditioning-induced mRNA translation patterns. We thus built a static gene regulatory network (GRN) using weighed gene co-expression network analysis (WGCNA)^34^, encompassing all TRAP fractions in our dataset (i.e., representing all conditions across all cell types), totaling 77 samples (Ext. Fig 7a). Our network identified 7 modules (Fig 4b; Ext. Fig 7b-d; Suppl Table 8), plus a “grey” module, containing non-correlated genes, which were excluded from posterior analysis. We filtered the most highly correlated gene pairs (dissimilarity index < 0.1) and constructed a visual representation of the GRN (Fig 4c). *Post-hoc* gene ontology analysis revealed that each module was associated with specific biological features.

**Figure 6.**
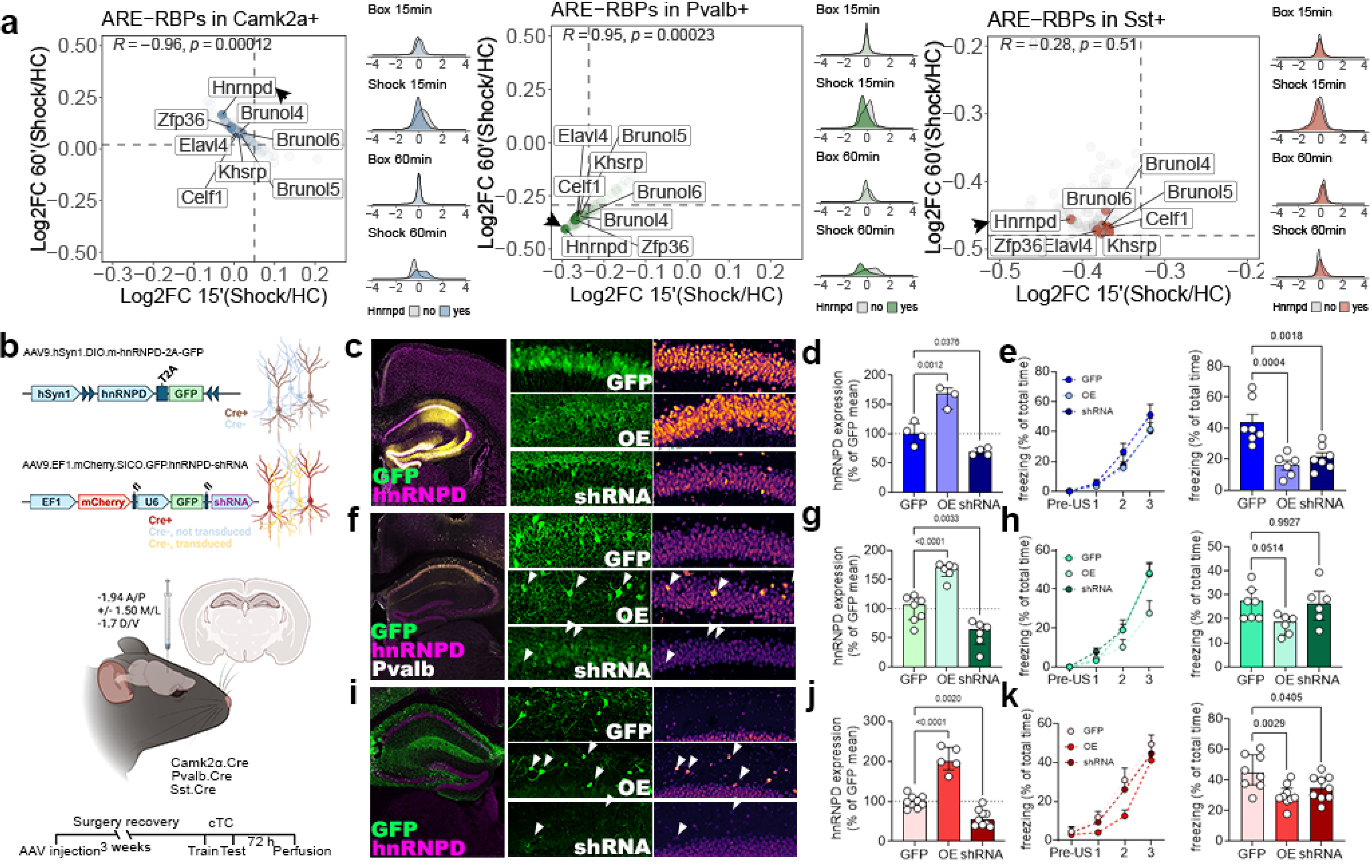
The neuron-specific disruption of ARE-based control of mRNA metabolism leads to memory malfunction. (a) Average fold changes of mRNAs predicted to be targets of ARE-RBPs in *Camk2a^+^, Pvalb^+^* and *Sst^+^* neurons (names of the RBPs are highlighted inside each plot). Top left corner = *R* represents Pearson’s correlation coefficient; *p* is the significance of the correlation. On the right of each plot, density plots show fold change distribution of targets of hnRNPD (colored) versus non-targets (in gray). Number of mice per group in the dataset used is listed in *Methods* section. (b) Schematics of bi-directional manipulation approach to target hnRNPD expression in a cell type-specific way. Green neurons = GFP^+^ neurons; red neurons = mCherry^+^ neurons; yellow neurons = GFP^+^/mCherry^+^ neurons. A/P = anterior/posterior. M/L = medial/lateral; D/V = dorsal/ventral coordinates. (c) On the left: representative images of the viral delivery to the dHPC. (d) Quantification of hnRNPD levels in AAV9.GFP-, OE- or shRNA-transduced neurons. Mean ± SEM. (e) Contextual threat conditioning of mice injected with AAV9.GFP-, OE- or shRNA viruses. On the left, training session; On the right, test session. OE = overexpression. Hab = habituation (N=6-7 mice per group); Mean ± SEM; *F* for panel d = 0.7307; *F* for panel e *=* 0.6853. (f-h) Same as in c-e, but for *Pvalb*^+^ neurons (N=6-7 mice per group); Mean ± SEM; *F* for panel g = 0.7307; *F* for panel h *=* 1.097. (i-k) Same as in c-e, but for *Sst*^+^ neurons (N=8-9 mice per group); Mean ± SEM; *F* for panel j = 0.0421; *F* for panel k *=* 1.412. *P* values are demonstrated to the fourth digit above bar graphs in each panel. One-Way ANOVA corrected with Tukey’s *post-hoc* test.

We next asked which modules were most associated with specific aspects (i.e. cell type, time point, or condition) of the dataset used to build the GRN. This revealed that the “yellow” and “brown” modules, which encompass mRNAs encoding RPs and mitochondrial proteins, respectively, had the highest correlation with the *shock* group (Fig 4d). In contrast, the “turquoise” module, containing mRNAs related to post-translation processing of proteins, showed a negative correlation with the *shock* group. Gene-trait correlation revealed that IEGs such as *Dusp5*, *Nr4a1* and *Junb* are among the best genes correlating with the *shock* group (Ext. Fig 7e). On the other hand, we found a suprising subset of genes with a high negative correlation with *shock*. For example, *Mettl18* has been described to methylate ribosomal proteins, thereby slowing translation elongation^35^ and modulating ribosomal biogenesis^36^; *Chd6*, a helicase involved in chromatin remodeling; and *Usp29*, a protein de-ubiquitinase. In addition, we found that “green” and “red” modules, which contain genes related to cellular respiration and RNA binding, had strong positive correlation with interneurons. In contrast, the “blue” and “black” modules, which gather genes related to RNA polymerase II-mediated transcription and trans-synaptic signaling, correlated mostly with excitatory neurons. Thus, the GRN had modular representation of genes highly correlated with specific aspects of our dataset.

WGCNA presents the advantage of finding global patterns of conditioning-induced translatome shifts that fit every cell type analyzed, but it fails to identify cell type-specific effects. To circumvent this problem, we mapped the log_2_FC values found in the differential expression analysis to the GRN (Fig 4e). We identified discrete patterns of altered translation across modules for each neuron type that evolved differently across time. Surprisingly, *Sst*^+^ mRNA translation patterns were more similar to what was seen in *Camk2a*^+^ cells rather than *Pvalb*^+^. However, this finding is consistent with multiple reports demonstrating that the physiological properties of *Sst*^+^ cells in the dHPC are very similar to those of excitatory neurons, such as a lower threshold for action potential generation^37^. In addition, we found that the turquoise module, originally presented as an inverse correlation with the *shock* group, showed a specific upregulation in excitatory neurons 60 min after conditioning (Fig 4e, heatmap). Gene ontology of the differentially expressed RA-mRNAs belonging to the turquoise module in excitatory neurons dataset (25% of the total module) revealed that these mRNAs encode proteins mostly related to proteostasis, such as ER homeostasis (e.g.*, Tmx1, Ptges3*, *Emc2, Emc7*), ubiquitinylation (e.g., *Ube* family of genes and *Rnf139*) and protein folding (e.g., *Vapa, Pdzd8, Dnaja1, Dnaja2* and *Dnajb4*). These findings point to a cell type-specific feature where excitatory neurons require tight control of proteostasis to cope with increased translation during memory consolidation^11,12^.

Given our findings that GADD34 regulated the conditioning-induced translation programs triggered in excitatory neurons, we hypothesized that its depletion would lead to loss of coordinated mRNA translation in these cells. To examine this, we mapped the fold changes found in the TRAP-seq of Camk2a.Cre-GADD34^flfl^ mice to the GRN (Fig 4f). Importantly, we found that while control mice displayed the same pattern as before, GADD34 knockout mice showed no activation of turquoise module. Instead, they displayed significant upregulation of the brown, green and yellow modules, related mostly to mRNA-protein interaction and mitochondrial metabolism. Altogether, our findings show that the GRN represents specific patterns of RA-mRNAs recruitment across neuron types and that this is disrupted in excitatory neurons in the absence of GADD34.

### Cell type-specific changes in RA-mRNAs induced by conditioning are controlled by abundance of regulatory elements present in the 3’UTR of mRNAs

mRNA translation is controlled by multiple mechanisms, including cis-(i.e. features present in the mRNA sequence/structure) or trans-regulatory elements (i.e. elements that target mRNAs or ribosomes and control translation). Cis elements were previously shown to control mRNA stability, localization and translation, and are often found in the 3’UTR^38^. We thus interrogated whether cis elements in mRNAs would dictate rules guiding conditioning-induced mRNA translation. We collected information regarding the composition of the following mRNA cis-elements from literature: m6A enrichment, number of AU-rich elements (AREs) in the 3’ UTR^39^, number of RBPs targeting the 3’ UTR, number of miRNAs targeting the transcript, and codon optimality^40^. We originally included 3’UTR length as a predictor, but its high correlation with both number of AREs and targeting miRNA suggested high levels of collinearity that may complicate interpretation of the model (Ext Fig 8-10, see correlograms). Therefore, we opted to exclude it from the final model.

To investigate whether mRNA regulatory elements were predictive of enrichment or depletion of an mRNA transcript in the translatome following conditioning, we leveraged classical machine learning techniques to build predictive models trained on the cis-regulatory features described above. Separate gradient-boosted decision tree models (XGBoost) were trained per cell-type and per condition to effectively capture the varied mRNA programs present in each condition. We assessed model performance by the area under curve (AUC) of their receiver operating characteristic (ROC) plots as well as accuracy metrics (Fig. 5b). We found that datasets reflecting changes at 60 min post-conditioning displayed the highest model performance (shock 60’ AUC: *Pvalb*^+^ = 0.94±.01, *Camk2a*^+^ = 0.95±.01, *Sst^+^* = 0.85±.05. Overall, this suggests that, regardless of the neuron type, the features embedded in the mRNA sequence drive ribosome loading after conditioning.

We asked whether the density of these mRNA features could explain the modular organization found in our GRN (Fig 5c). Importantly, the features were unevenly distributed across modules, and *black* and *turquoise* modules displayed significant enrichment of all features (except codon optimality) when compared to the other modules (Fig 5d, statistical analyses are in Suppl Table 9). We matched the feature density with the fold changes found in each experimental group of our dataset (we excluded *box* controls here, as their XGBoost models did not perform as well as *shock* groups). The heatmap denoted an inverse correlation between the time point of upregulation of RA-mRNAs and the density of features found in the mRNAs in excitatory neurons (Fig 5c, compare bottom panel with upper panel). For example, the turquoise module, highly expressed in excitatory neurons at the 60 min timepoint, displayed the highest concentration of features. On the other hand, the brown module, increased at 15 min in the same cells, had much lower feature accumulation. Interestingly, this analysis further revealed that gene modules displaying accumulation of these features had poor translation performance in *Pvalb*^+^ neurons. Instead, these neurons displayed coordinated expression of genes with less amounts of regulatory features. Finally, we did not observe any clear pattern that could explain the regulation of translation in *Sst*^+^ cells during consolidation.

To quantify the relationship between the accumulation of regulatory features and differential expression of RA-mRNAs, we analyzed the covariation between the number of features and the log_2_FC values occurring in each condition (Fig 5e). In this analysis, it became clear that AREs have an inverse relationship with conditioning-induced changes in RA-mRNAs across multiple modules, particularly in interneurons (see second row in each heatmap split). Furthermore, we found that AREs directly correlated with higher RA-mRNA fold changes in excitatory neurons at the 60 min timepoint. This led us to hypothesize that AREs have a crucial role in explaining the differential modulation of RA-mRNAs during consolidation. Thus, we performed permutation importance tests on each XGBoost model (Fig 5f, Ext Fig 8-10). We performed this analysis only in *shock* 60 min datasets, as these were the best performing models. We found that, for *Camk2a^+^* and *Pvalb^+^* neurons, permuting the number of AREs consistently resulted in the greatest decreases in validation accuracy. This was not the case for *Sst*^+^, where no feature had clear distinct importance for model performance. This result indicates that ARE count was the most predictive feature of up or downregulation for *Camk2a^+^* and *Pvalb^+^* models. This finding was confirmed again when XGBoost models trained on single features from all shock 60’ datasets performed well when trained on the ARE feature set (ARE-shock 60’ AUC: *Pvalb*^+^ = 0.90, *Camk2a*^+^ = 0.92, *Sst*^+^ = 0.70 (Ext Fig 11). These results suggest that mRNA regulatory elements determine the pool of RA-mRNAs associated with memory consolidation. Additionally, the results show that AREs are the principally informative features for our classical machine learning approach, suggesting that AREs play a distinct role in differential mRNA translation following conditioning.

### The neuron-specific disruption of ARE-based control of mRNA metabolism leads to memory malfunction

AREs are targeted by numerous RBPs that can promote stabilization, translation or decay of the targeted mRNA^41^. These RBPs can be present in different cell types with different stoichiometries, changing the patterns of ARE-dependent RNA metabolism. We examined the expression patterns of 10 different ARE-targeting RBPs in the translatome of *home cage* and found differences across neuron types (Ext. Fig 12a). For example, Celf-encoding mRNAs are significantly enriched in excitatory neurons. On the other hand, *Elavl2*, *Elavl4* and *Hnrnpd* were enriched in *Pvalb*^+^ neurons. Furthermore, we found that the expression of many of these RBPs is affected by conditioning (Ext. Fig 12b). For example, *Hnrnpd* is significantly upregulated in excitatory neurons after training, whereas it is downregulated in *Pvalb*^+^ cells. Overall, these data provide initial evidence that ARE-dependent mRNA metabolism is significantly altered follow conditioning in different neuron types.

We proceeded to examine the role RBPs played in the training-induced translatome modifications. We used RBPMap^42^ to map motifs at the 3’UTR that were targeted by 126 different RBPs (Suppl. Table 10). All the mRNAs targeted by a given RBP had their log_2_FC values transformed into arithmetic means that were then used as coordinates for centroid positioning on a two-dimensional plot (Ext. Fig 13a). We found a notable correlation in centroid positioning when we compared shock 60’ and shock 15’ groups (R*_Pvalb_* = 0.98; R_Camk2a_ = -0.94; R_Sst_ ^=^ -0.28, Ext. Fig 13b; Suppl. Table 11). The box control group displayed good correlation values (R*_Pvalb_* = -0.72; R_Camk2a_ = 0.45; R_Sst_ ^=^ -0.75), but the effects were milder and not significant when compared to the centroid for all genes (Ext Fig 13). To confirm that these results were reliable, we also collected data from the literature that experimentally defined mRNAs targeted by RBPs. We collected data from 47 different RBPs, 14 of which overlapped with the prediction assay. Overall, similar analyses demonstrated the same correlational trends, thus reinforcing the reliability of our findings (Ext Fig 13).

Due to our finding that AREs are the best predictors of conditioning-induced differential translation of mRNAs, we restricted our analyses to RBPs that target AREs. We found that their targeted mRNAs were downregulated in *Pvalb*^+^ and upregulated in *Camk2a*^+^ neurons 60 min following conditioning (Fig 6a). Surprisingly, we found that mRNAs targeted by ARE-targeting RBPs were downregulated in *Sst^+^* neurons, an apparent conflict with results obtained above, where number of AREs do not have high predictive value in the *Sst*^+^ model. We posit that, rather than total counts of AREs, the role of AREs in regulating the changes to the translatome in *Sst*^+^ cells rely on expression patterns of ARE-targeting RBPs, which will require further investigation. The modulation seemed to be ubiquitous to all ARE-targeting RBPs, consistent with the notion that these proteins compete with each other for their target mRNAs. Our analysis indicated that heterogeneous nuclear Ribonucleoprotein D (hnRNPD) displayed significant changes in mRNA translation of its client mRNAs across datasets (Suppl. Table 11), suggesting a universal role for this RBP in memory consolidation. hnRNPD is primarily found in the nucleus, participating in alternative splicing^41^. It is also known to destabilize mRNAs via de-adenylation, resulting in decay^43^, thereby granting it a major role in the regulation of the mRNA pool present in the cell. We hypothesized that its expression in neurons is essential for proper long-term memory formation. We perturbed expression of hnRNPD by either overexpressing or silencing it in each neuron type studied (Fig 6b). We found the most notable role for hnRNPD in Camk2a^+^ and *Sst^+^* cells, where either increasing or decreasing its expression caused memory impairments (Fig 6c-e; Fig 6i-k). In *Pvalb*^+^ cells, we found that increasing, but not decreasing, hnRNPD expression caused memory impairment (Fig 6f-h), in accordance with expression data harvested from the translatome (Ext. Fig 12b). Altogether, these findings demonstrate a clear, ubiquitous role for hnRNPD in the regulation of memory formation and reinforce the power of the dataset we have generated to be a highly effective tool to aid in the discovery of new molecular correlates and modulators of memory.

## Discussion

The creation and stabilization of long-term memories require orchestrated communication between cells in different brain regions, as well as the local tuning of activity through the interaction of neighbor neurons in microcircuits. Thus, each neuron type presents molecular and electrophysiological signatures that enable them to effectively play their specific role during memory consolidation. This molecular framework dictates how neurons dynamically respond to external cues, such as converging network stimuli, causing adaptations that are specific to each neuron type. Here we probed the alterations to the translational programs induced by threat conditioning in three different neuron types: *Camk2a^+^, Pvalb^+^* and *Sst^+^*. Using the translatome as a readout of molecular programs triggered during consolidation proved to be valuable, as it focuses on mRNAs bound in ribosomes that are likely undergoing active translation. This approach offers key advantages over other commonly used strategies to study neuronal plasticity at the molecular level, as the translatome more effectively approximates the proteome—the functional outcome of translation that ultimately drives plasticity dynamics. To our knowledge, this is the first comprehensive report of distinct neuron type-specific translational programs of the dHPC during memory consolidation. Our results demonstrate that the initial pattern of cellular adaptation is diverse, both in the nature of the cellular processes/structures altered and in the molecular mechanisms that guide the translation of the proper set of mRNAs.

One of the most noticeable features in the datasets that we collected was that *Camk2a*^+^ neurons displayed a dynamic response within the first hour of consolidation, promoting translation plasticity as early as 15 minutes after conditioning by increasing the production of controllers of protein metabolism. This rapid shift in the production of proteostasis machinery is noteworthy, given that it is comparable to the speed of translation of immediate early genes^44^. Interneurons displayed a delayed proteostatic reconfiguration, instead prioritizing rapid changes in regulators of neurotransmission. One possible interpretation of such a phenomenon is that negative feedback systems tend to be tightly controlled, such as to produce phasic inhibition of increasing excitatory activity in the hippocampus^45^. Thus, it is possible that interneurons, after targeting proteins that control synaptic transmission, then respond with a ubiquitous reconfiguration of protein metabolism.

The changes in the nature of mRNAs being translated by neurons during consolidation are accompanied by an increase in translation, dependent on the dephosphorylation of the initiation factor eIF2α that is enabled by the scaffolding protein GADD34^8,11,12,24,27,46^. GADD34, traditionally considered to be a stress-related effector, promotes protein phosphatase 1-induced dephosphorylation eIF2α after acute cellular stress is resolved, restoring protein synthesis to its homeostatic levels^26^. However, in neurons, we found that GADD34 is repurposed to regulate rapid increases in protein synthesis, regardless of the presence of a stressful stimulus. We suggest that GADD34 mediates the capacity of neurons for large increases in *de novo* translation, promoting translation of mRNAs related to protein folding, degradation and translation initiation. This finding is unexpected, given the canonical role assigned for GADD34 under stressful conditions^25^. Based on our findings, we hypothesize that the repurposing of GADD34 is necessary for neuronal resilience, not only promoting traditional neuronal plasticity, but also ensuring proper cellular adaptation to a stressful condition that accompanies plasticity. Indeed, pathways associated with buffering cellular stress, such as the unfolded protein response^47^, chaperone production^48^ and alternative translation initiation pathways^49^, are consistently linked to neuronal plasticity and memory formation. Their impairment, on the other hand, plays a fundamental role in synaptic and cognitive failures observed in neurodegenerative diseases^50–52^. Overall, these findings reinforce the notion that neurons simultaneously engage both a large-scale anabolic response and a stress-resolving pathway during plasticity, a previously undescribed cellular state that warrants further investigation.

In addition to regulating increases in mRNA translation via upstream signaling molecules, such as GADD34, mRNAs carry cis-regulatory elements in their sequence that are recognized by the cell machinery, which ultimately decides the fate of that mRNA. These elements were previously shown to be crucially involved in subcellular localization of mRNAs in neurons^38^, and to be involved in the translational control of specific mRNAs^53^. Here, we have demonstrated that cis-regulatory elements have a critical role in regulating mRNA dynamics during long-term memory consolidation. The model employed here is currently limited by knowledge of static features pertaining to the mRNA sequence. This can be solved via a deep characterization of cis- and trans-features occurring in specific types of neurons, which will provide insights into how the molecular diversity of neurons determines their role during memory consolidation. Our data suggests that translation of mRNAs during consolidation is tightly linked to both the nature and number of cis-regulatory elements present within each mRNA molecule. Furthermore, we identified and characterized a previously unknown modulator of long-term memory formation, hnRNPD. This protein is associated with post-transcriptional mRNA decay^43^ and alternative splicing^41^, and its microdeletion results in intellectual disability in humans^54^. It remains to be determined how hnRNPD, with its two main post-transcriptional roles in mRNA modulation, controls memory formation. Nevertheless, approaches such as those we have deployed in the current work will enable the identification of numerous additional proteins that play key roles in cognitive tasks and neurological disease.

In summary, we have examined the changes in the translatome induced by threat conditioning in three different neuron types known to be involved in memory consolidation. Our results indicate a clear pattern of response by each neuron, distinguishable by specific biological features rapidly modified in an adaptive response to afferent stimuli. However, we find that the translatome modifications are governed by elements hard coded in the mRNA sequence, regardless of the cell type examined. These universal rules governing cis-elements may then interact with different molecular environments across cell types to produce distinct outcomes. Our data represent a powerful hypothesis-generating resource for the scientific community, as our novel findings reinforce the uniqueness of the translatome as an invaluable tool to understand neuronal plasticity and the molecular mechanisms of memory. Overall, we anticipate this study to benchmark the usage of the translatome and the investigation of RNA metabolism dynamics as a framework to dissect neuronal plasticity at the cellular and systems levels. We believe that future studies utilizing our dataset and this approach will have broad impact in the collective fields of molecular, systems, and computational neuroscience.

### Limitations of the study

Due to current technical limitations, it is not feasible at present to simultaneously study the transcriptome and the translatome of different neuron types in high throughput settings. Due to this limitation, we cannot rule out that a fraction of dynamic shifts in the translatome may occur due to transcriptional modulation. We note, however, that even though this might hold true, it does not alter the interpretation of the findings gathered here. Further, we highlight that TRAP utilizes the overexpression of a single ribosomal protein, RPL10a, which can selectively increase translation of specific transcripts via internal ribosome entry sites^55^. These ribosomes were precluded from analyses and may contain additional important information on how the translatome responds to threat conditioning. Finally, this work reflects the average modifications that occur in different neuron types, not accounting for intrapopulation variability. This was by design, and the reasons were two-fold: (1) single-cell analyses of the translatome in dynamic settings such as the one examined here remain an unresolved challenge; (2) due to the shallow nature of scRNA-sequencing, single-cell analyses hinder the examination of nuanced molecular modifications, requiring the incorporation of pseudo-bulk analyses to reach meaningful conclusions, ultimately leading to similar analyses as the ones performed here. However, we cannot exclude that some of the features found in our datasets are restricted to neuronal sub-populations of the dHPC, a possibility that remains unexplored.

## Supporting information

Supplemental data supporting the main manuscript

## Acknowledgements.

We thank the Genome Technology Center from New York University School of Medicine for the excellent RNA library preparation and RNA sequencing involved in this project, particularly Gael Westby. We thank Dr. Arkady Khoutorsky for the generous donation of AAV9.Camk2a.RPL10a::eGFP virus, used in Figure 2. This study was funded by NIH, grant NS 122316.

## Author contributions

M.M.O. and E.K. conceptualized and designed the project. M.M.O. and R.C. performed bioinformatics and computational analyses. M.M.O. collected and processed all TRAP samples. M.M.O., O.M., W.J.L., E.M., K.S.A.R., C.L., C.S. and E.H.L. performed and analyzed experiments. E.K. and T.C. contributed animals, materials and analysis tools.

## Methods

### Mice

All animal protocols were reviewed and approved by the New York University Animal Care and Use Committee (protocol # 2021-1139). Mice were provided with food and water *ad libitum* and maintained in a 12h-12h light-dark cycle at New York University at a stable temperature (78oF) and humidity (∼50%). Male and female mice were equally distributed across experimental groups and used between the ages of 3-5 months. B6;129S6-Tg(Camk2a-cre/ERT2)1Aibs/J (Camk2a.Cre^ERT2^), B6.129P2-Pvalb^tm1^(cre)^Arbr^/J (Pvalb.Cre), Sst^tm2.1^(cre)^Zjh^/J (Sst.IRES.Cre), B6;129S4-Gt(ROSA)26Sor^tm9^(EGFP/Rpl10a)^Amc^/J (TRAP^fl^) and B6.Cg-Tg(Camk2a-cre)T29-1Stl/J (T29.Cre) mouse lines were obtained from Jackson laboratories. B6.129P2-Ppp1r15a^tm1.1Ajf^/Mmnc (GADD34^fl^) was obtained from Mutant Mouse Resource & Research Centers (MMRRC). Every mouse strain was backcrossed with C57Bl/6J mice from Jackson laboratories every 5 weeks, to ensure minimal background variation. T29.Cre, Pvalb.Cre or Sst.Cre were crossed with TRAP^fl^ mice to obtain Camk2a.Cre-TRAP, Pvalb.Cre-TRAP or Sst.Cre-TRAP mouse lines. Heterozygous GADD34^fl/+^-Heterozygous Camk2a.Cre^ERT2^ were crossed with heterozygous GADD34^fl/+^ mice to obtain Camk2a.CreERT2.GADD34^fl/fl^. Heterozygous GADD34^fl/+^-Homozygous Pvalb.Cre were crossed with heterozygous GADD34^fl/+^ mice to obtain Pvalb.Cre.GADD34^fl/fl^. Heterozygous GADD34^fl/+^-Homozygous Sst.IRES.Cre were crossed with heterozygous GADD34^fl/+^ mice to obtain Sst.IRES.Cre.GADD34^fl/fl^.

### Behavior

All behavior was conducted during the light cycle. Mice were randomly assigned for the order of testing. All data were collected by experimenters’ blind to mouse genotype. To habituate mice, over the course of two days mice were placed in the threat conditioning room for 1hr then each mouse was handled for 2min each day. For simple contextual threat conditioning, mice were placed in the context, which consisted of a metal rod floor and white house light, first for 298s and then presented with three series 2s, 0.6mA foot shocks. The first and second inter-trial intervals were 128s and 98s respectively, and after the last shock, mice remained in the chamber for an additional 170s. This training protocol lasted in total 700s. LTM was tested 24h after training by placing mice back in the same context for 300s. Freezing behavior was automatically measured using Freeze Frame 4 (ActiMetrics).

### Translating ribosome affinity purification (TRAP)

TRAP was performed as before^15,24^. In brief, we separate the procedure into three steps: (1) anti-GFP-coated magnetic bead preparation; (2) sample collection and ribosome purification; (3) RNA extraction. All steps were performed in RNAse-free condition. Total number of mice per experimental group is listed below:

*Camk2a* 15 min dataset

. Home cage = 3 mice

. Box only = 3 mice

. Shock = 4 mice

*Camk2a* 60 min dataset

. Home cage = 4 mice

. Box only = 4 mice

. Shock = 5 mice

*Pvalb* 15 min dataset

. Home cage = 4 mice

. Box only = 4 mice

. Shock = 5 mice

*Pvalb* 60 min dataset

. Home cage = 6 mice

. Box only = 4 mice

. Shock = 6 mice

*Sst* 15 min dataset

. Home cage = 4 mice

. Box only = 3 mice

. Shock = 5 mice

*Sst* 60 min dataset

. Home cage = 6 mice

. Box only = 5 mice

. Shock = 6 mice

Below we list each step of the TRAP purification process:

### Anti-GFP-coated magnetic bead preparation

300 μl of MyOne Streptavidin beads per reaction was collected, washed once with PBS, and incubated with 120 μl of recombinant protein L suspension (1 μg/ml) and 180 μl of PBS for 35 minutes/RT/rotating. Beads were washed 5x with 1 ml PBS + 3% IgG-free bovine serum albumin solution and incubated with anti 50 μg of each anti-GFP Htz antibodies for 1h/RT/rotating. Beads were then washed 3x with cold low-salt buffer (80 mM HEPES-KOH, 600 mM KCl, 40 mM MgCl_2_, 1% NP-40, 1x Halt protease/phosphatase inhibitor, 0.5 mM DTT, 100 ug/ml Cycloheximide) and resuspended in 200 μl low-salt buffer per reaction. Beads were stored at 4oC until time of use.

### Sample collection

After contextual threat conditioning training session, mice were returned to their home cage for 15 minutes or 60 minutes. Mice were quickly euthanized via cervical dislocation, and the brain removed and rapidly transferred to ice-cold PBS + 100 μg/ml Cycloheximide. The dorsal hippocampus was micro-dissected and transferred to an ice-cold pestle container containing 1 ml of tissue lysis buffer (80 mM HEPES KOH, 600 mM KCl, 40 mM MgCl2, 1% NP-40, 1x Halt protease/phosphatase inhibitor, 0.5 mM DTT, 100 μg/ml Cycloheximide, RNAsin). Tissue was homogenized with 12 strokes of a motorized pestle and the suspension transferred to an Eppendorf tube. The homogenate was cleared via centrifugation (2000 *g*/10 min/4°C) and the supernatant saved. The supernatant was added with 1/9^th^ v:v of 300 mM DHPC, mixed by inversion and incubated for 5 minutes on ice. Samples were then centrifuged at 20.000 *g/*10 min/4°C and supernatant collected. 1/10^th^ of the volume was saved for total lysate sequencing, and the remaining volume was added with 200 μl of anti-GFP-coated magnetic beads and incubated overnight/rotating/4°C. On the next day, samples were briefly spun to recover all beads and washed 4x with 1ml ice-cold high-salt buffer (80 mM HEPES-KOH, 1.4M KCl, 40 mM MgCl2, 1% NP-40, 1x Halt protease/phosphatase inhibitor, 0.5 mM DTT, 100 μg/ml Cycloheximide, RNAsin). Beads were then resuspended in 350 μl of room temperature RLT buffer added with 10% β-mercaptoethanol (RNEasy plus kit). 350 ul of RLT buffer + BME was further added to total lysate samples.

### RNA purification

RNA purification was done using RNEasy plus kit (Qiagen). This kit was chosen due to the extra purification step to eliminate gDNA contamination. RNA extraction was performed in accordance with the suggested protocol, with following changes: (1) 50 μl of elution buffer was used, to maximize recovery; (2) two sequential elution steps were performed. RNA quality was measured using Agilent’s ScreenTape for global assessment of RIN, and for detailed quantitative and qualitative assessment Agilent’s Bioanalyzer was used.

#### Library preparation and sequencing

The SMART-Seq HT Kit (Takara) was utilized to generate complementary DNA (cDNA) from 1 ng of total RNA input. The synthesis process included 13 PCR cycles. The quality and concentration of the stock RNA were assessed using an Agilent Pico Chip on the Bioanalyzer system, ensuring high-quality RNA inputs. cDNA was quantified using the Invitrogen Quant-iT system and subsequently diluted to a concentration of 0.3 ng/µL for library preparation. Libraries were prepared using the Nextera XT DNA Library Preparation Kit, employing a total of 12 PCR cycles to amplify and uniquely barcode the samples. The quality of the generated libraries was assessed using the Agilent High Sensitivity TapeStation to verify fragment sizes and integrity. Additionally, the concentration of libraries was measured with the Invitrogen Quant-iT system to ensure accurate normalization prior to pooling. Libraries were pooled in equimolar ratios and sequenced on the Illumina NovaSeq X+ platform using paired-end 100-cycle reads. The sequencing data was processed using standard bioinformatics pipelines.

#### Alignment to the genome and feature count

Sequencing adapters and low-quality bases were trimmed using *Cutadapt* version 1.8.2 using the following arguments (-q 20 -O 1 -a CTGTCTCTTATA)^57^. Trimmed reads were then aligned to the mm9 mouse genome assembly using STAR version 2.7.7a and raw gene counts were obtained using *-quantmode* argument^58^. Total feature counts were obtained using *featureCounts*^59^. Feature counts were extracted and converted into one single raw count matrix, where each row was a gene, and each column was a sample. When samples were obtained from different cohorts of mice, we used ComBat-Seq^60^ to batch-correct.

#### Bioinformatics

All analyses described below were performed in R 4.4.2, except the XGBoost classification model, which was done in Python, using *scikit-learn* package. All codes are available upon request or downloadable from GitHub.

#### Quality control and filtering of enriched genes

The datasets were subjected to a two-stage filtering process and data clearing to ensure strict data analysis. One of the noticeable features across datasets was the increase in translation of mRNAs traditionally associated with glial cells when mice were conditioned, such as markers *Cx3cr1* and *Aldh1l1*. Because these are known to be low or not expressed by neurons, we decided to filter our datasets to select truly neuron-enriched transcripts in the translatome. To achieve this in an unbiased manner, we used the following approach to filter out genes that were not “truly enriched” in the purified fraction:

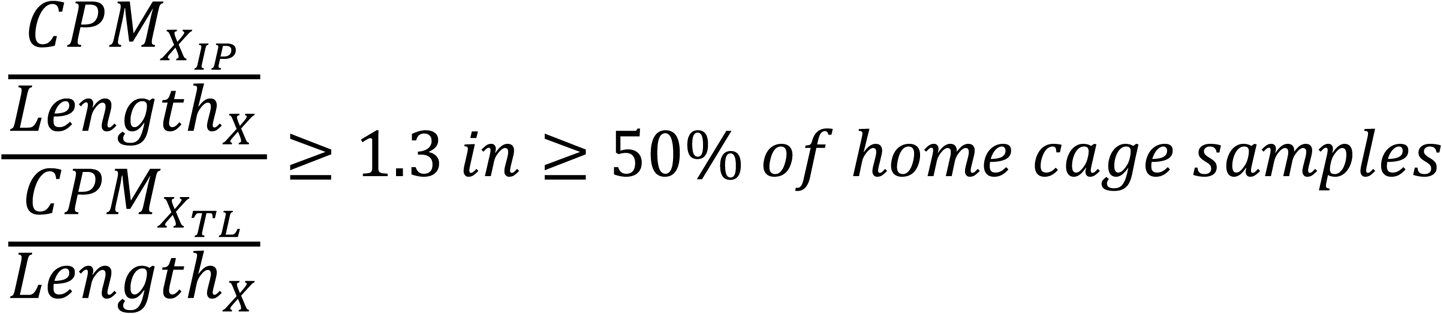

Where 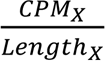 is the Reads per Kilobase Million (RPKM) per gene, IP is the counts in the purified fraction, and TL is the counts in the total fraction (*total lysate*). Overall, the equation states that, when a gene has a 1.3-fold increase in RPKM values in the purified fraction versus total lysate, it is considered enriched. To increase stringency and avoid inherent problems that come with sequencing datasets - such as noisy expression in a fraction of genes – we included an extra step, where the gene should be enriched in at least 50% of the samples of the baseline (or *home cage* group). Here, we did not include the experimental groups, as they may shift the expression patterns of RA-mRNA expression, resulting in condition-specific enrichment of translating mRNAs. Instead, these conditions were filtered to have the same gene list obtained in *home cage* samples. We ended with gene lists enriched in a cell-specific way that represented 40-60% of the total genes originally detected in the sequencing. The filtering process was performed for each independent neuron type, yielding cell-specific enriched gene list.

Following the initial filtering, we next excluded samples that did not present enrichment of neuron type-specific markers. We considered enrichment as a 2-fold increase in RPKM values of TRAP versus total lysate fractions. On average, we excluded 1-2 samples per dataset using this threshold. The cell type markers used were:

*Camk2a^+^ neurons = Camk2a, Slc17a6, Slc17a7*

*Pvalb^+^ neurons = Gad1, Pvalb*

*Sst^+^ neurons = Gad1, Sst*

*Glial markers* (to evaluate depletion) *= Aldh1l1, Gfap, Cnp, Cx3cr1*

### Differential expression analysis and Gene Ontology analysis

Identification of DEGs was made using DESeq2 package, from R ^61^. Briefly, crude count matrices were batch corrected using ComBat-seq^60^. The resulting matrix served as input to DEseq2^61^. Transcripts with less than 20 counts in more than half of the samples were filtered out. Datasets were then normalized by the size of library, and variance stabilized using the *vst*() function from DEseq2 package. DEG identification was done by filtering genes that had *p* adjusted value < 0.05 in comparisons between conditions. After a full list of DEGs was obtained, genes categorized as enriched per cell type (see above) were filtered in and kept for future analysis. We opted for doing the filtering at this later stage to ensure the rigor of adjusting the *P* value in the whole list of genes. For intra-cellular analyses, genes identified as enriched in that particular cell type were kept. For inter-cellular analyses, such as the WGCNA (see below), a full list, containing all genes enriched in all neuron types, was obtained and used as input. A complete list of all DEGs from all datasets is available in Supplementary table 3. These lists of DEGs were independently subjected to Gene Ontology analysis using the *clusterProfiler* package from R, using false discovery rate (FDR) as *post-hoc* correction method and only sets larger than 30 genes were considered in the analyses^62^.

### Weighted gene co-expression network analysis (WGCNA)

WGCNA was performed using the *WGCNA* R package ^34^. Briefly, a variance-stabilized (vst) data frame containing all conditions from all cell types was fed into the pipeline, as suggested in the original package vignette. For *Camk2a* 15 min time point, we found a significant reduction in the total amount of represented genes (∼1000 less genes than in other datasets). We thus imputed those values using classification and regression trees method, from the *mice* R package, in the pre-vst dataset. No noticeable difference in the raw count distribution was observed, suggesting that the imputation did not significantly alter the nature of our dataset. To design a scale-free network, the premise behind WGCNA, we first calculated the power for soft-thresholding, which uses lasso regularization to select significant connections in the network. A power of 20 was found to reach the threshold of 0.8 in the scale-free index, suggested by the designers of WGCNA as optimal for producing scale-free networks.

The categorization of genes in co-expression modules was done using the *blockwiseModules()* function, using power = 20 and minimum module size of 30. We generated a signed network, to separate genes that had positive versus negative correlations. 10 modules were identified and assigned colors to them. The *grey* module, however, contained genes that were not assigned to any particular module, and was therefore excluded from functional analyses below. The gene ontology (GO) analysis for each module identified was performed using the *anRichment* package in R, with a threshold of 0.05 and Bonferroni correction. We manually curated GO terms per module to display in representative figure in panel 2c.

To design a network that could be exported and visualized in cytoscape^63^, we first generated a weighted network using the function *TOMsimilarityFromExpr()* from *WGCNA* package. We then exported all the most significant connections (threshold = 0.1) from all modules but *“grey*” using the *exportNetworkToCytoscape()* function. The edges file was imported into cytoscape and matched with the nodes file, containing the information regarding gene-module membership. Modules were colored in the network with colors matching their names. To map log2FC values obtained in the differential expression analysis into the network, we matched the log2FC of each gene with the corresponding network node, and mapped both color (upregulated = *red*, downregulated = *blue*) and size (below threshold = *size 5*, above threshold = *size 30*). Genes that were in the third quartile of log_2_FC (i.e. 25% of most differentially expressed genes) were considered above threshold.

### Prediction of RBP targets

3’UTR from all transcripts identified in each sequencing were retrieved using biomaRt. The longest variant of 3’UTR was selected by transcript. The 3’UTRs were submitted as query to RBPMap^42^, using the following parameters: *-db_motifs ‘all’ -stringency ‘high’ -conservation ‘off’*. Results were extracted into a data frame using scripts built *in house* (Supplementary table 7). For centroid plots, the arithmetic means of all mRNAs predicted as targets to a given RBP were calculated for each time point and plotted in a two-dimensional space.

### Machine learning model

In order to begin our machine learning analysis, we constructed cell-type specific datasets consisting of 7 features of transcripts identified from our TRAP-seq analysis: AU-rich elements (ARE), codon optimality (tAI), UTR length, total number of known regulatory RNA binding proteins (RBPs), total counts of m6A modification, total number of known regulatory microRNAs (miRNA), and mean FPKM. These feature sets were obtained from different sources:

> . ARE counts in the 3’ UTR were retrieved from previous work that quantified them using regular expression in the longest transcript version^39^.
>
> .Codon optimality was retrieved from http://stadium.pmrc.re.kr/^40^. Codon optimality is represented as the geometric means of the relative optimality of codons from the coding sequence of each mRNA.
>
> . UTR lengths were retrieved using biomaRt database (R).
>
> . Total number of known regulatory RNA binding proteins were retrieved using scripts built *in house* to identify motifs in the longest version of the 3’UTR of each transcript that are targeted by RBPs included in the RBPMap^42^ database (see above).
>
> . total counts of m6A modifications per transcript, per cell type, were obtained from a study that generated a single-cell database of the epitranscriptome^64^.
>
> . total number of regulatory microRNAs were obtained for each cell type. Specifically, we first retrieved a list of microRNAs expressed in each neuron type from previous literature^65^. We then used miRWalk^66^ to identify microRNA-binding regions in the longest isoform of the 3’UTR of each transcript identified per neuron type. Finally, we used scripts built *in house* to filter the binding sites to keep only the ones targeted by microRNAs expressed in the specific neuron type.
>
> . mean FPKM values were derived from the translatome dataset of *home cage* mice and represent the average ribosomal occupancy of each mRNA in resting conditions.

Furthermore, the dataset was filtered to select most notable fold changes relative to *home cage* (log2FC > 0.5 or log2FC < -0.5). Those transcripts which were deemed to be significantly up or downregulated in comparison to home-cage mice were used for training. This selection thereby narrows the scope of the subsequent analysis as identifying those features of mRNAs that predict the regulatory valence of transcripts known to be differentially modulated during threat conditioning. Using this dataset, gradient-boosted decision tree ensemble models (XGBoost v2.0.3) were trained using 5-fold cross validation (CV). Per CV fold, feature columns were scaled to ensure they were distributed such that µ = 0 and σ = 1. After scaling, outlier detection was performed using the IQR with bounds determined by the standard multiplicative factor 1.5, with values determined to be outliers removed and subsequently imputed using KNN imputation with K = 5 (scikit-learn v1.2.1). Hyperparameters were tuned using a Bayesian optimization strategy (scikit-optimize v0.9.0) by scoring optimal estimators using the accuracy of the trained estimator on the held-out validation folds. Due to the static nature of mRNA sequence regulatory features, optimal models were trained and selected per condition and per cell type, allowing for cross condition/cell type comparison. Optimal model topologies across conditions were compared using the mean area under the receiver operating characteristic (AUC-ROC) curve as computed per fold using the held-out validation data. Feature importances were determined using two methods. Firstly, untrained models with optimal hyperparameters were trained on single features of the dataset and performance across features was compared using the AUC computed per fold using the held-out validation data. Additionally, permutation importance tests were performed using 10 permutation sets per feature and importance was assessed by the mean reduction of model accuracy over these sets.

#### Viral injection

Mice aged 10-12 weeks were anesthetized with isoflurane (induction: 5%, maintenance: 2% in O2 vol/vol, via nose cone) and placed in a stereotaxic frame. Bilateral dorsal hippocampus (−1.94 mm AP, ±1.50 mm ML, and −1.5 mm DV) was stereotactically targeted with a 10ul Hamilton syringe. The syringe was lowered to the 0.2mm pocket below the target site and remained there for 5 min before moving to the target and beginning the injection. After the injection, the needle stayed in the target site for 5 min before it was withdrawn. A StereoDrive microinjector (Microinjection Robot, StereoDrive) was used to deliver 500 nl of viral solution (2.1-2.8 vg/ml titer) at a rate of 0.2ul/min. *Camk2a*.Cre, *Pvalb*.Cre, and *Sst*.Cre transgenic mouse lines were injected with either AAV9.hSyn1.DIO.m-hnRNPD.2A.eGFP or AAV9.EF1.mCherry.DIO.U6.GFP.hnRNPD-shRNA (Vector Biolabs). *Camk2a.Cre^ERT^*^2^ were also injected with AAV9.CBA.DIO.GADD34-mCherry. For all injections, AAV9.hSyn1.DIO.GFP was used as exogenous protein expression control. *Camk2a.Cre^ERT^*^2^*/GADD34^flfl^* mice were injected with AAV9.CAG.FLEX.RPL10a::eGFP (Addgene). Mice were given 3 mg kg−1 ketoprofen for 3 days post-op as an analgesic after surgery and allowed to recover for at least 2 weeks before behavioral experiments.

#### Retro-orbital administration of AHA

AHA and RO injections were performed as previously described^21^. Briefly, azidohomoalanine (AHA) (Vector Laboratories, CCT-1106) was dissolved in PBS at 25mg/ml and delivered at 50mg/kg to awake mice via retroorbital (RO) injection immediately after conditioning. RO injection was performed by two researchers, with the first restraining the mouse’s head before subsequently drawing back the skin below the eye and second researcher using a 27G needle to inject into the retro-bulbar sinus. Two hours later, brain tissue were collected as below.

#### Perfusions and brain sectioning

Mice were anesthetized with isoflurane and perfused trans-cardially with PBS to clear blood and then with 4% PFA in PBS (∼40 ml per mouse). Brain was removed and post-fixed in 4% PFA in PBS overnight/4°C. Brains were mounted in 3% agarose and sectioned using a vibratome Leica VT1200S. Around 40 sections of 40 μm thickness containing the dorsal hippocampus area were collected and kept free-floating in PBS added with sodium azide.

#### FUNCAT Staining

Sections were permeabilized with PBS + 0.5% Triton X-100 solution (15min/RT/agitation). Sections were then blocked in 5% normal goat serum (NGS) + 5% bovine serum albumin (BSA) + PBS + 0.1% Triton X-100 solution (60 min/RT/agitation). Sections were then incubated in Click reaction solution (Vector labs) to label AHA-containing peptides (overnight/4°C/rocking), following manufacturer’s recommendations, using a final concentration of 2.5 μM alkyne-647. Sections were washed 2x with 4 ml of PBS + 0.1M EDTA (10 min/RT/agitation), followed by two washes with 4 ml of PBS + 0.1% Triton X-100 (10 min/RT/agitation). After another blocking step with 5% NGS + PBS + 0.1% Triton X-100 (60 min/RT/agitation), sections were incubated with anti-NeuN primary antibody diluted in blocking buffer (1:500, overnight/4°C/rocking). Brain sections were then washed 3x PBS + 0.1% Triton X-100 (10 min/RT/agitation) and incubated with goat anti-guinea pig secondary antibody (AlexaFluor 488) diluted 1:500 in blocking buffer (90 min/RT/rocking). Finally, sections were washed 3x PBS + 0.1% Triton X-100 (10 min/RT/agitation), stained with DAPI solution (1 μg/ml in PBS) for 5 min/RT/agitation and mounted on glass slides using Prolong Antifade reagent.

#### Immunohistochemistry and imaging quantification

Sections were permeabilized with PBS + 0.5% Triton X-100 solution (15min/RT/agitation). Sections were then blocked in 5% normal goat serum (NGS) + PBS + 0.1% Triton X-100 solution (60 min/RT/agitation). Brain sections were then transferred to blocking buffer added with primary antibodies (for concentrations, please refer to antibodies and reagents section) and incubated overnight/4oC/agitation. They were subsequently washed 3x with PBS + 0.1% Triton X-100 (10 min/agitation/RT) and incubated with blocking buffer added with secondary antibodies (1:500, 90 min/agitation/RT). Sections were washed 3x again with PBS + 0.1% Triton X-100, transferred to PBS and mounted in glass slides. After sections were dry, Prolong was added to the slides and covered with a rectangular coverslip. Imaging was performed in an upright Leica SP8 system, with 20x magnification objective. To measure the efficacy of hnRNPD overexpression and silencing systems, we imaged 4 CA1-containing fields per mouse (2 sections, 2 images per section, one CA1-containing image per hippocampus). Quantification of hnRNPD intensity was done using Fiji macros built in house. Briefly, cell type-specific markers were used to generate a mask. That mask was used to select the region-of-interest in the channel containing hnRNPD staining. The raw integrated density value was obtained and normalized against the total area selected. For better visualization in the graph (Fig 5c-e), GFP-injected signal was averaged, and the resulting value was determined as 100%. All the points were then re-calculated to obtain a relative value against the average of control.

#### RNAscope

RNAscope coupled with immunohistochemistry was performed following manufacturer’s standard procedure, using anti-GFP (Rb, 1:300) for counter-staining. No modifications to the protocol were done, except skipping antigen retrieval step. Probe against *Ppp1r15a* was purchased on C1, *Camk2a* and *Pvalb* on C3, and *Sst* on C4.

#### Tamoxifen preparation and administration

Tamoxifen was prepared and administered as suggested by Jackson laboratories. In brief, tamoxifen powder was diluted in corn oil to the final concentration of 20 mg/ml, under vortex agitation for 4h/RT. Solution was stored at 4°C for up to a week. Mice were administered with tamoxifen solution intraperitoneally for 5 days, 100 μl per day (∼ 75 mg/kg).

#### Statistical analysis

Mice were included as independent biological variables in all the experiments performed in this work. No replicated measures were obtained from the same mouse and treated as independent samples. Graphs were designed using *ggplot2* package from R or GraphPad Prism. The exact test and post-test corrections are described in figure legends, but as a general rule we used two-tailed unpaired *t* test for comparisons between two groups, and One-Way ANOVA for comparisons that involved more than one group, with all conditions independent from one another. One-Way ANOVA was corrected using Tukey *post-hoc* test. Prior to statistical analysis, data was tested for normality using Shapiro-Wilk test, and Q-Q and residual plots used for visual inspection of normality.

