## Supplemental data supporting the main manuscript for "Neuron type-specific mRNA translation programs provide a gateway for memory consolidation"

[#]Co-second authors

**Supplementary file**

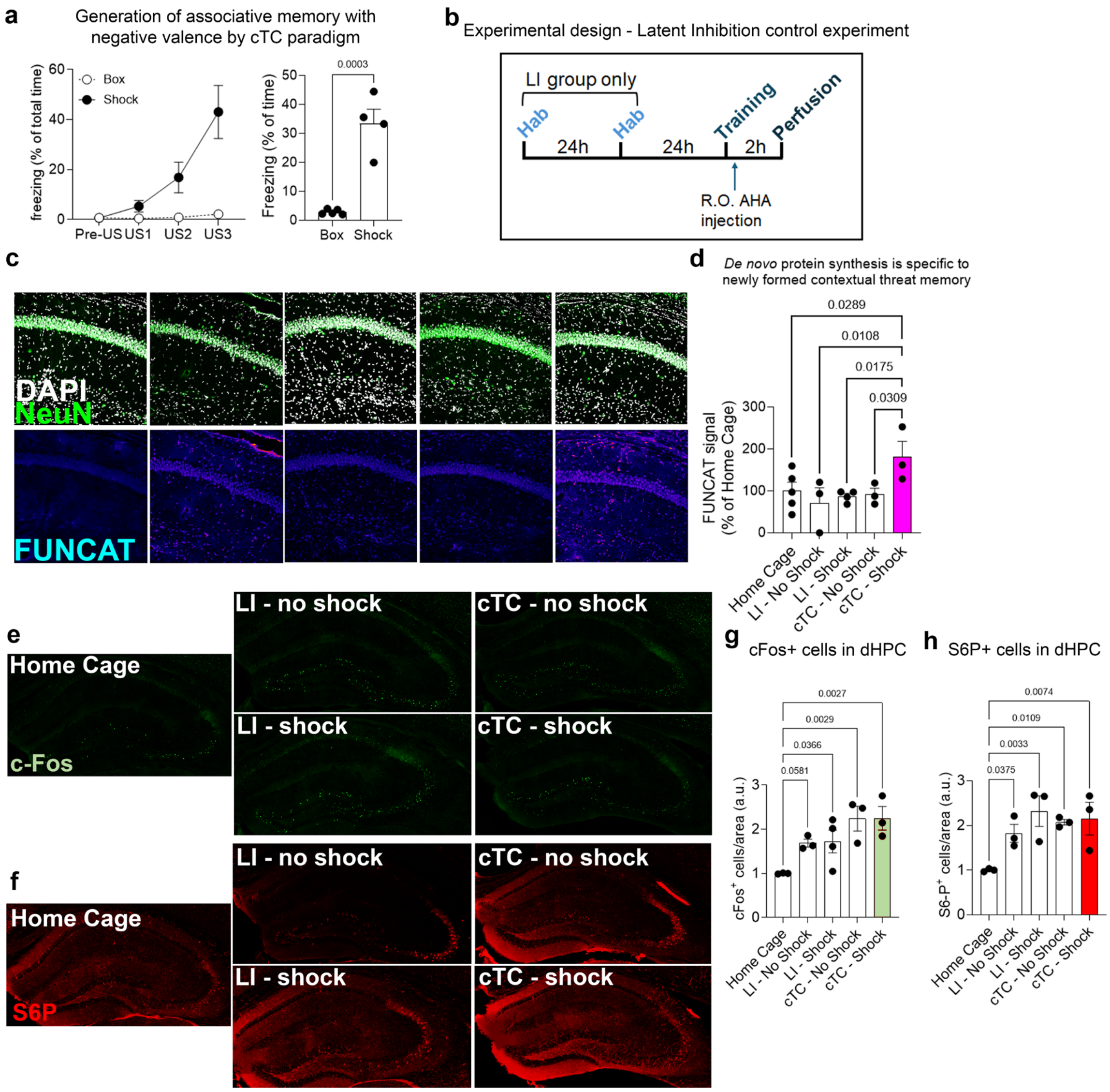


**Ext. Fig 1. Contextual threat conditioning is a robust method to study consolidation-associated neuronal *de novo* protein synthesis. (a)** Validation of the contextual threat conditioning method employed in this study. Left panel: training session. US = Unconditioned stimulus; Pre-US: period prior to first shock delivery. Each point represents the average of freezing displayed inter-trial. Right panel: Test session, held 24h after training session. Box = Presentation to new context without shock; Shock = Presentation to new context with shock. Student's *t* test; number represents the *p* value. **(b)** Experimental design for the latent inhibition experiment. Latent inhibition (LI) group was presented to the contextual information during 15 min sessions, spaced 24h. During this period, mice were allowed to freely explore the area. On the training session, LI or naïve mice were presented to the context, and shocked or not. As baseline controls, *home cage* mice were used. Retro-orbital injection of AHA was delivered immediately after the training sessions. Mice were kept for 2h in their home cage and perfused with 4% PFA. N = 3-5 mice per group. **(c)** Representative images of FUNCAT staining in mice that underwent the LI experiment. Top panels: White = DAPI; Green = NeuN; Bottom panels: FUNCAT. **(d)** Quantification of (c). **(e)** c-Fos staining (green) in the dHPC of mice that underwent the LI experiment. **(f)** S6P staining (red) in the dHPC of mice that underwent the LI experiment. **(g)** Quantification of (e). **(h)** Quantification of (f). One Way ANOVA with Tukey *post-hoc* correction. Numbers represent the *p* values of specific comparisons. Mean ± SEM.

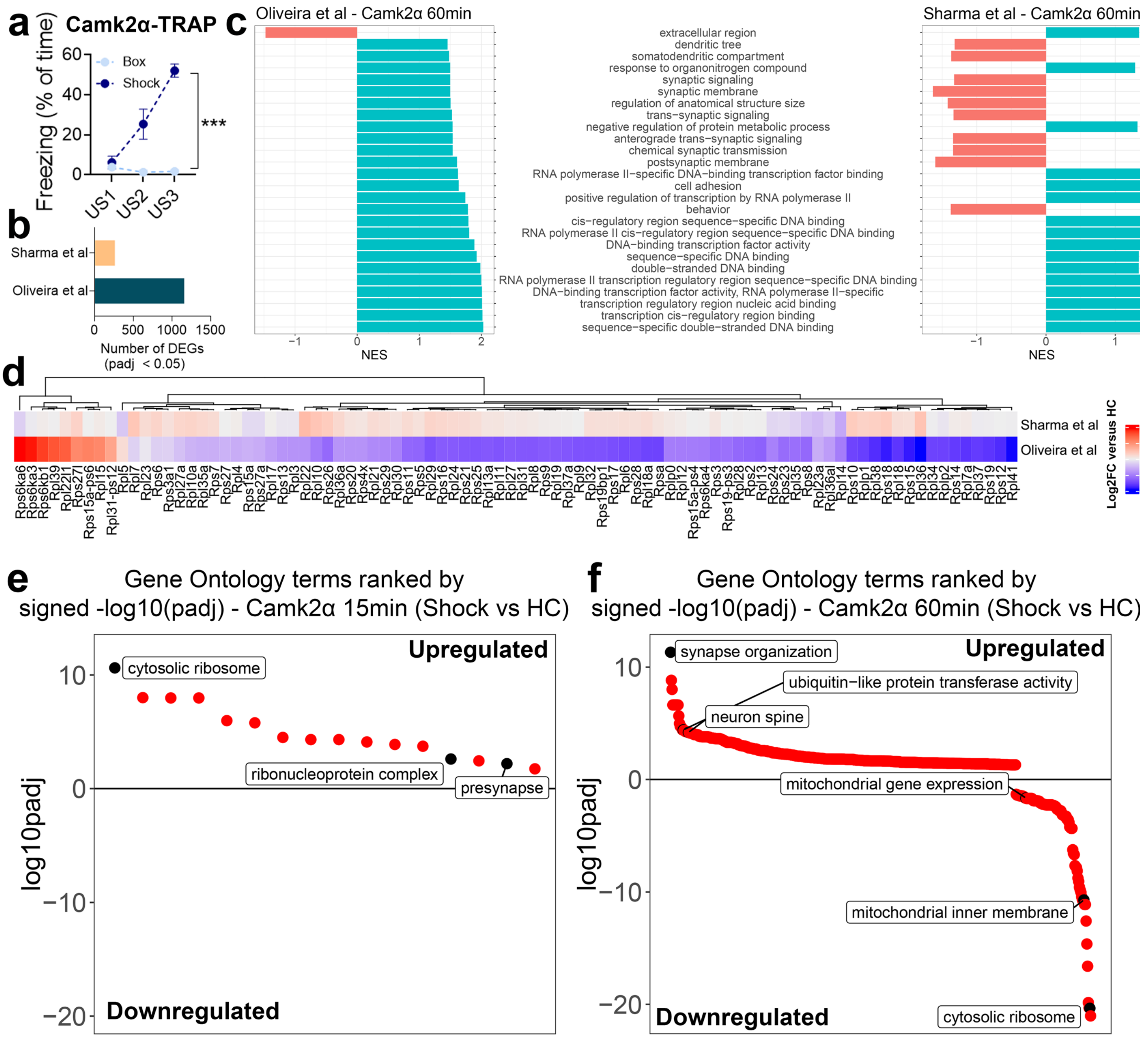


**Ext. Fig 2. Characterization of molecular programs triggered in excitatory neurons in response to threat conditioning. (a)** Freezing time (represented as % of total time) of mice used in the TRAP experiments during training session (N = 3-5 mice per group). Light blue = *Box* control; Dark blue = *Shock*. One-Way ANOVA repeated measures; *** = $p < 0.0001$. Mean ± SEM. **(b)** Comparison of absolute number of differentially expressed genes (DEGs; *p adj* < 0.05) found in our dataset versus what was found by Sharma and colleagues. **(c)** Comparative gene ontology analysis of DEGs found in our work versus Sharma and colleagues. Dark cyan = Upregulated terms; Light red = downregulated terms. The GO terms are specified between plots. NES = Normalized Enrichment Score. **(d)** Heatmap representing Log2FC of ribosomal proteins in the dataset collected in this work versus Sharma and colleagues. Colors represent standardized Log2FC. Blue = downregulated; red = upregulated. **(e)** Ranked dot plot of all terms found in the gene ontology analysis of DEGs identified 15 min after threat conditioning in *Camk2a*$^{+}$ neurons. GO terms are ranked according to *p adj.* Terms above the horizontal line are upregulated, below are downregulated. Black dots: GOs represented in the main manuscript. **(f)** Same as (e), but for the 60 min time point.

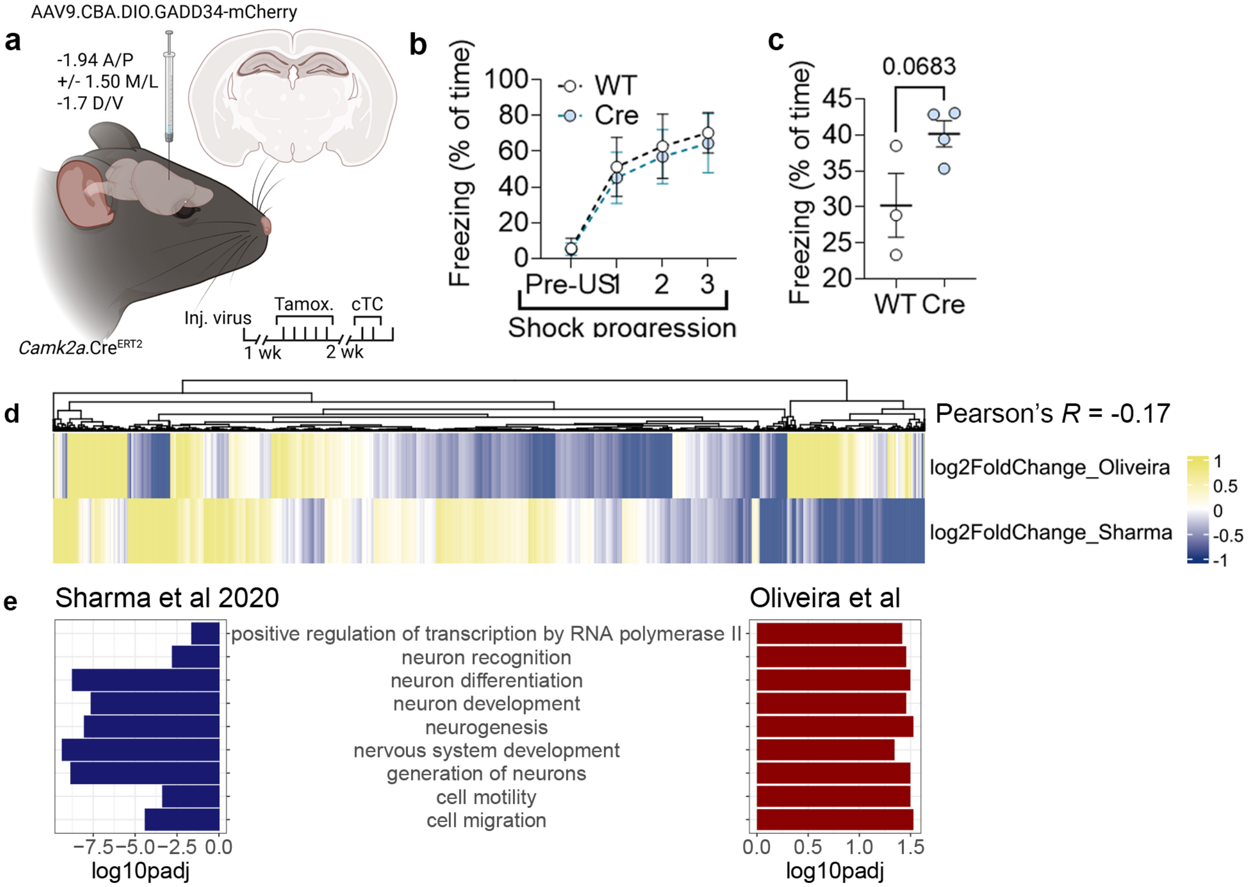


**Ext. Fig 3. GADD34 regulates molecular programs associated with long-term memory consolidation.** **(a)** Illustration representing the viral injection strategy and experimental design. AAV9.CBA.DIO.GADD34-mCherry was injected in the dHPC and mice daily administered with tamoxifen 1 week after surgery, for 5 days. Contextual threat conditioning was performed two weeks after the last tamoxifen injection. **(b)** Training session of contextual threat conditioning of mice injected with AAV9.CBA.DIO.GADD34-mCherry (N=3-4 mice per group). US = Unconditioned stimulus; Pre-US: period prior to first shock delivery. Each point represents the average of freezing displayed inter-trial. Mean ± SEM. **(c)** Testing session of contextual threat conditioning (N=3-4 mice per group). Student's *t* test, number on panel represents *p value*. Mean ± SEM. **(d)** Heatmap demonstrating the differences between DEGs found in $GADD34^{fl/fl}$ mice (higher eIF2$\alpha$-P) and eIF2$\alpha^{S51A/A}$ mice (lower eIF2$\alpha$). Note the low Pearson's *R* correlation on top right, suggesting the two manipulations do not cause opposing effects on the translatome, as we would expect. **(e)** Gene ontology analyses of both studies identified nine terms regulated in opposing ways by the two mutations. The terms names are listed between the bar graphs. On the left, results from Sharma and colleagues, showing negative regulation; On the right, the findings from this manuscript, showing positive regulation of these terms.

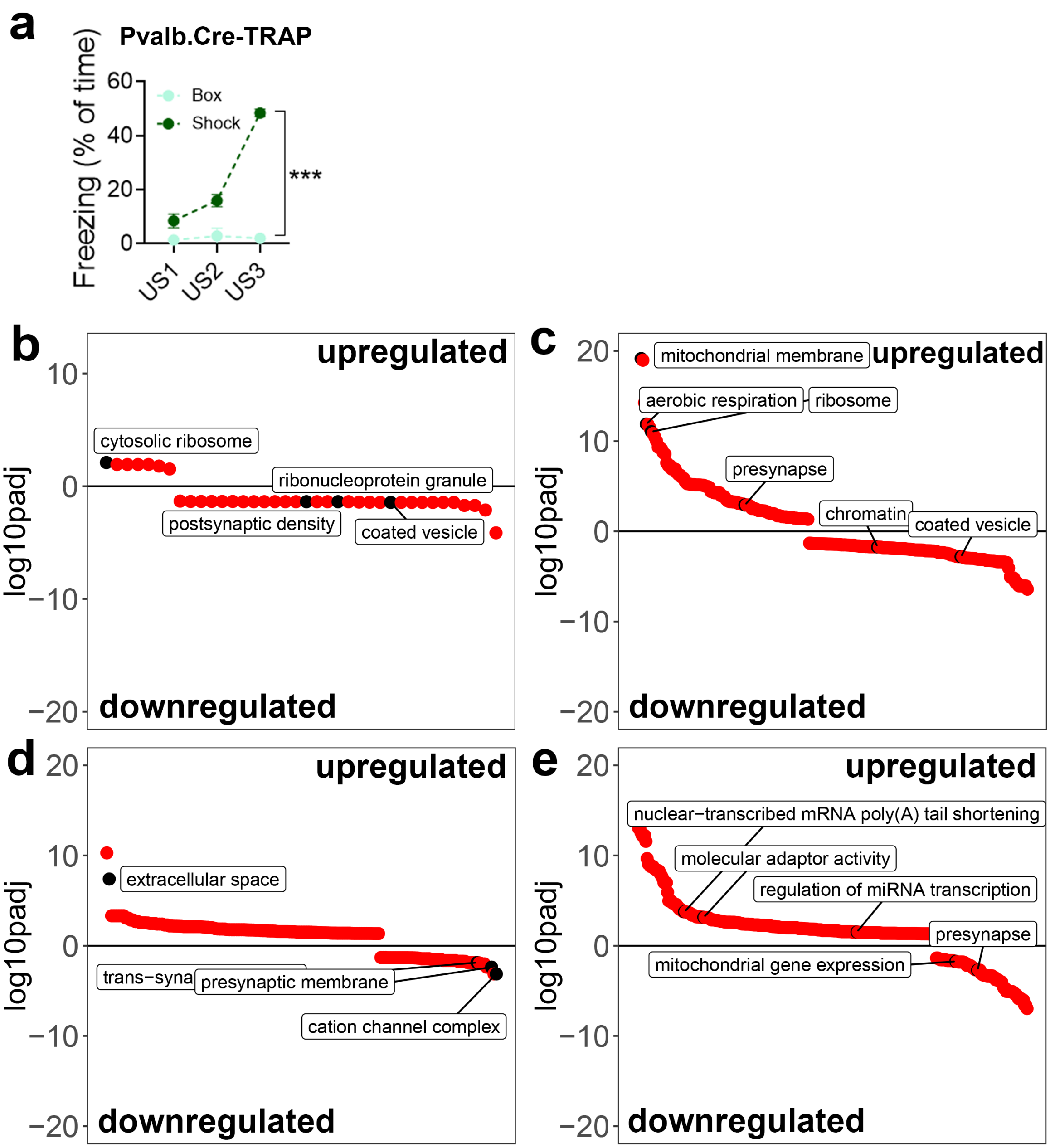


**Ext. Fig 4. Characterization of molecular programs triggered by interneurons during memory consolidation. (a)** Freezing time (represented as % of total time) of mice used in the TRAP experiments during training session (N = 4-6 mice per group). Light green = *Box* control for *Pvalb.*Cre mice; Dark green = *Shock* group for *Pvalb*.Cre mice; Light red = *Box* control for *Sst.*Cre mice; Dark red = *Shock* group for *Sst.*Cre mice. US = Unconditioned stimulus (shock). One-Way ANOVA repeated measures; *** = $p < 0.0001$. Mean ± SEM. **(b-e)** Ranked dot plot of all terms found in the gene ontology analysis of DEGs identified after threat conditioning in *Pvalb*$^{+}$ and *Sst*$^{+}$ neurons. GO terms are ranked according to *p adj.* Terms above the horizontal line are upregulated, below are downregulated. Black dots: GOs represented in the main manuscript. **(b)** *Pvalb*.Cre 15 min time point; **(c)** *Pvalb*.Cre 60 min time point; **(d)** *Sst.Cre* 15 min time point. **(e)** *Sst.Cre* 60 min time point.

**Oliveira et al.,**

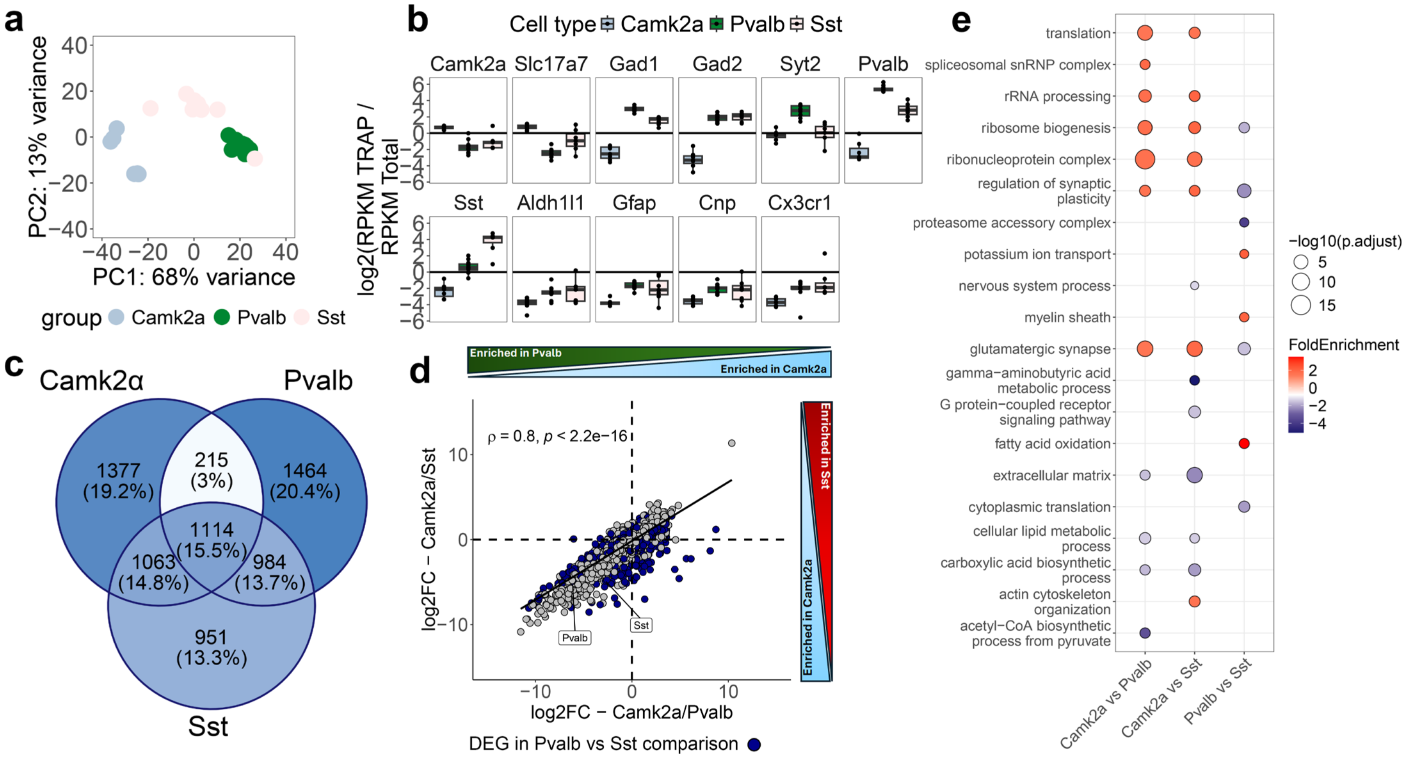


**Ext. Fig 5. Characterization of cell type-specific translatomes and their differences. (a)** Principal Component Analysis (PCA) of baseline translatomes (*home cage* mice). Each dot represents an individual mouse. dark green = *Pvalb*+; Light-green = *Sst*+. *N* = 6-9 mice/group. **(b)** Enrichment of cell type-specific markers in the purified translatome of each cell type. *Camk2a* and *Slc17a7* = excitatory cell markers; *Gad1, Pvalb, Sst* = Inhibitory cell markers; *Gfap, Cnp, Cx3cr1* = Glial markers. The upper and lower whiskers represent the largest and smallest value within 1.5 times above or below the interquartile range, respectively. **(c)** Venn diagram of the mRNAs identified in each cell type-specific purification. Color intensity illustrates the size of the set. **(d)** Spearman correlation of Log2FC of *Camk2a*$^{+}$ resting translatome versus *Pvalb*$^{+}$ (x axis) or *Sst*$^{+}$ (y axis). Spearman Rho = 0.8. Blue dots = mRNAs differentially modulated in the *Pvalb* vs *Sst* comparison. **(e)** GO analysis of DEGs found in the comparison between resting cell type-specific translatomes. Yellow means enriched in the first cell type in the comparison, magenta means enriched in the second cell type in the comparison. Color is relative to fold enrichment; size is mapped to significance (-log$_{10}$ *p adj*).

**Oliveira et al.,**

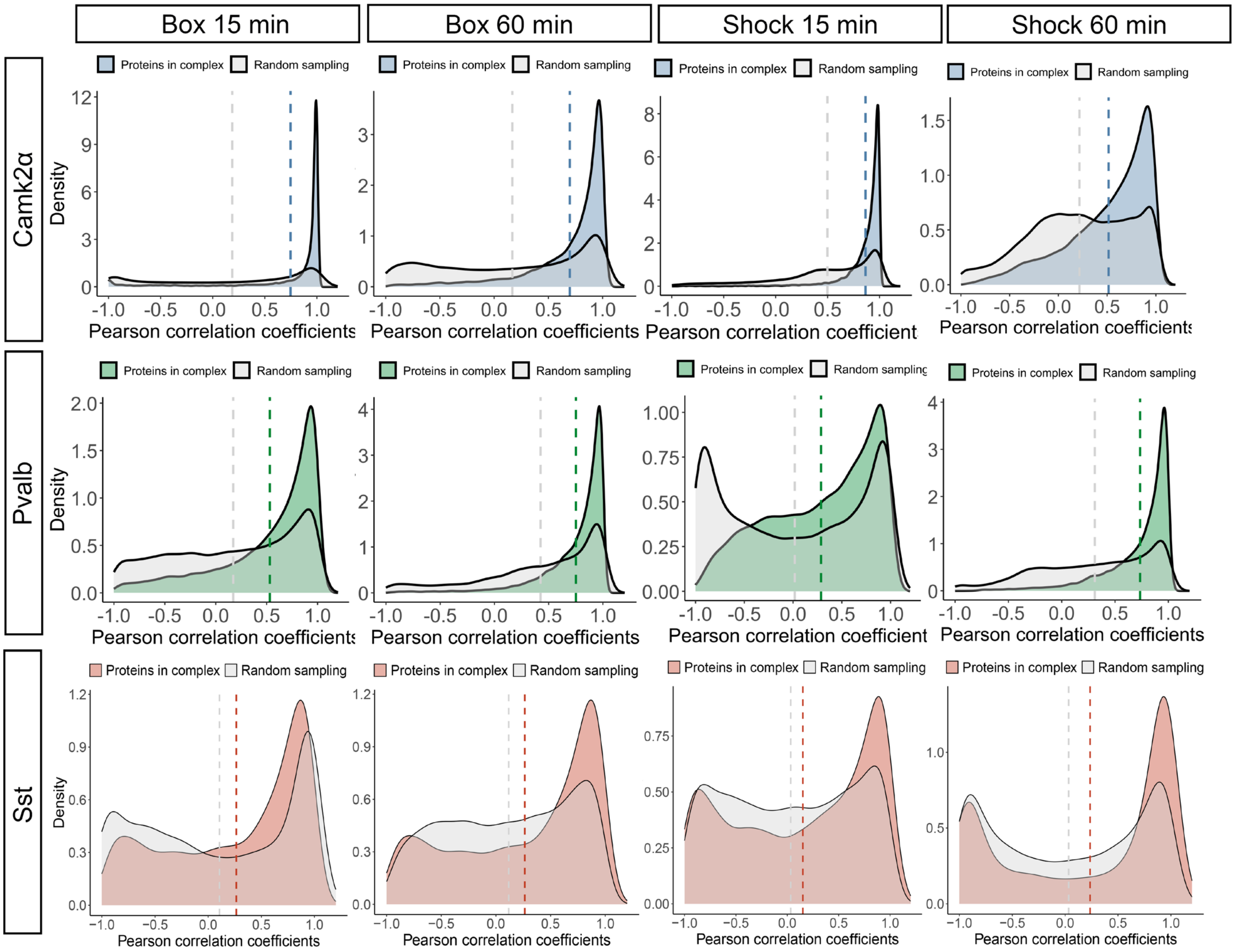


**Ext. Fig 6. mRNAs encoding proteins belonging to the same functional unit have higher correlation of expression than non-related mRNAs across cell types.** Gene-gene pairwise correlation coefficients of expression patterns were calculated for mRNAs encoding proteins belonging to the same protein complex. As a control, gene pairs were randomly sampled from the dataset and their correlation coefficients calculated. Dashed vertical lines indicate the mean of Pearson's correlation coefficient for each group. Note how the correlation of expression patterns is more similar among mRNAs belonging to the same functional units. Blue graphs = *Camk2a*; Green graphs = *Pvalb*; Red graphs = *Sst.*

Oliveira et al.,

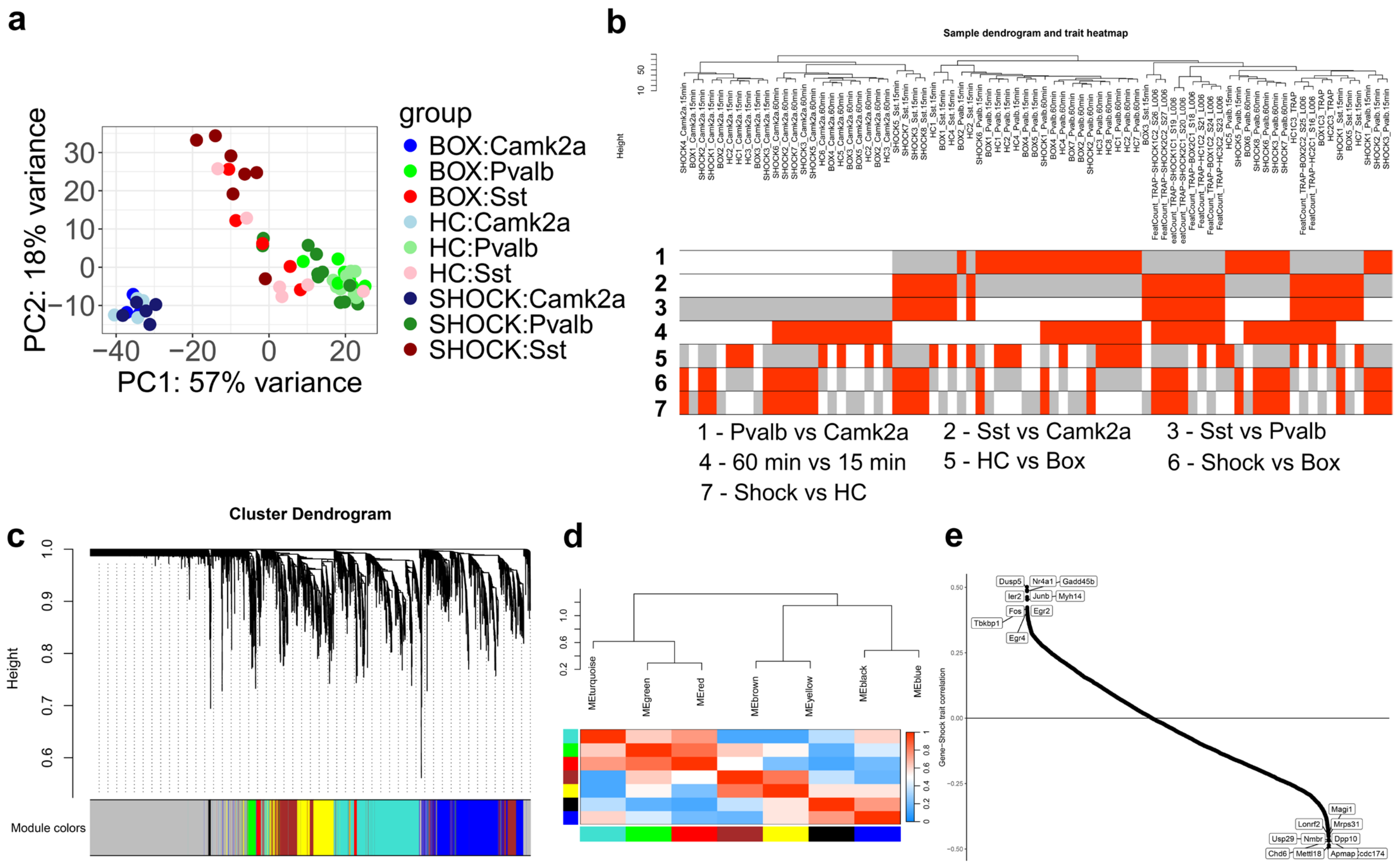


**Ext. Fig 7. Validation of the scale-free network generated using WGCNA.** (a) Principal component analysis of the whole dataset included in the WGCNA. (b) Dendrogram representing the clustering of samples by similarity. The rows under the dendrogram represent the relationship of each sample with external traits used: 1-3 comparisons between cell types; 4 – comparisons between time points; 5-7 – comparisons between time points. (c) Dendrogram and linear color map representing the clustering of genes per module. Colors represent the name of the module identified. (d) Cross-correlation of the 10 modules generated by WGCNA. The gray module was not included, as it represents genes that were not inserted in any cluster. Red = high correlations; Blue = low correlations. (e) Rank plot ordered by correlation of each gene against the *Shock* trait. Each dot represents a gene. Highlighted are the 10 genes with highest direct or 10 highest indirect correlation against the trait. Note the presence of multiple IEGs in the ones with highest trait correlation.

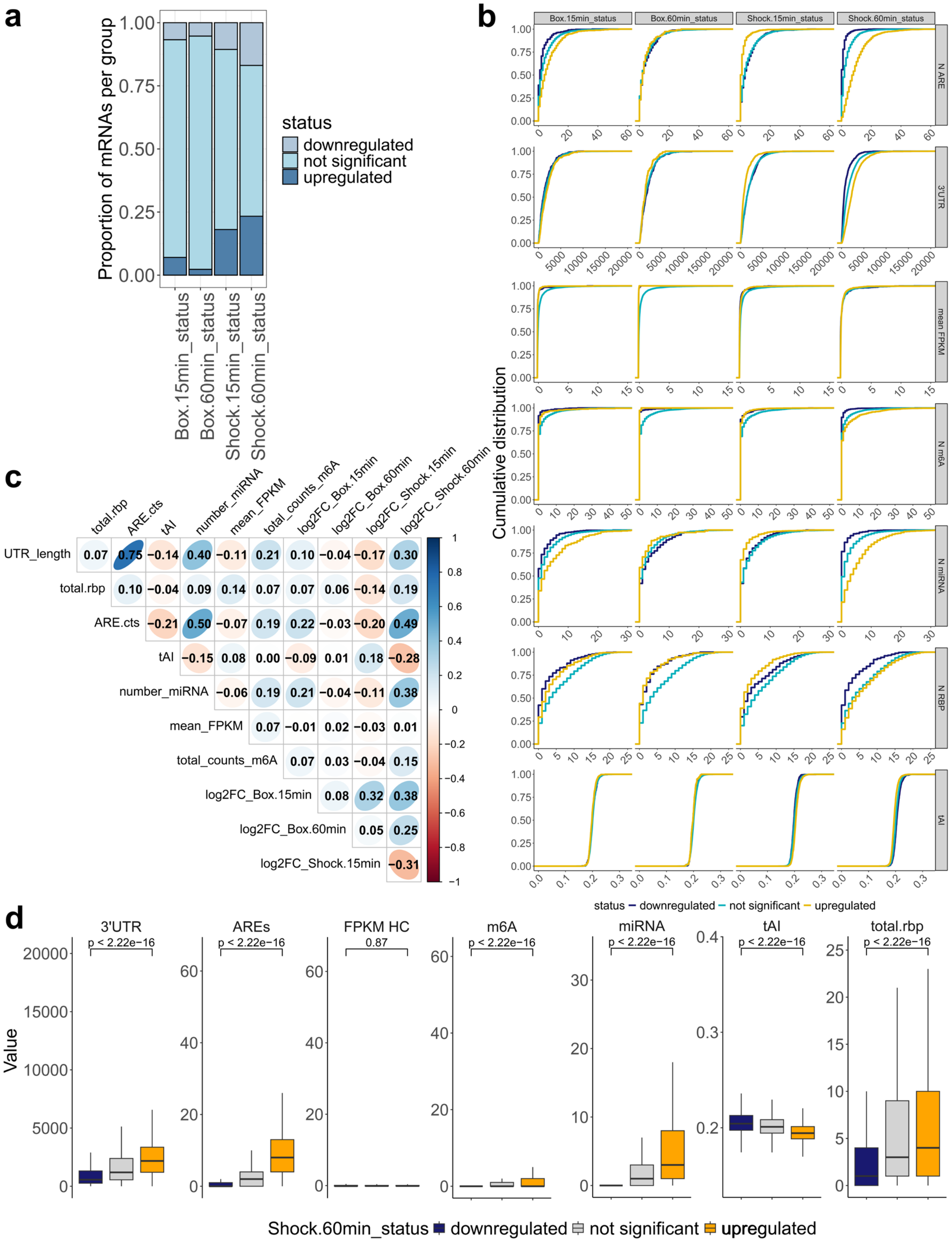


**Ext. Fig 8. Characterization of the input dataset for the *Camk2a*⁺ ML model.** (a) Stacked bar plot illustrating the proportion of genes defined as *downregulated* (top fraction), *not significant* (middle fraction) or *upregulated* (bottom fraction) per group modeled. (b) Cumulative distribution function of each mRNA element, per group modeled. Blue = *downregulated*; Light blue = *not significant*; Orange = *upregulated*. (c) Correlogram illustrating the linear relationship between fold change values of each dataset (normalized to *home cage* controls) and mRNA elements. Numbers inside each square represent the R coefficient. Blue = positive correlation; Red = negative correlation. (d) Box plots directly comparing the prevalence of each mRNA element in *downregulated* versus *upregulated* genes. One-Way ANOVA corrected with Tukey's *post-hoc* test.

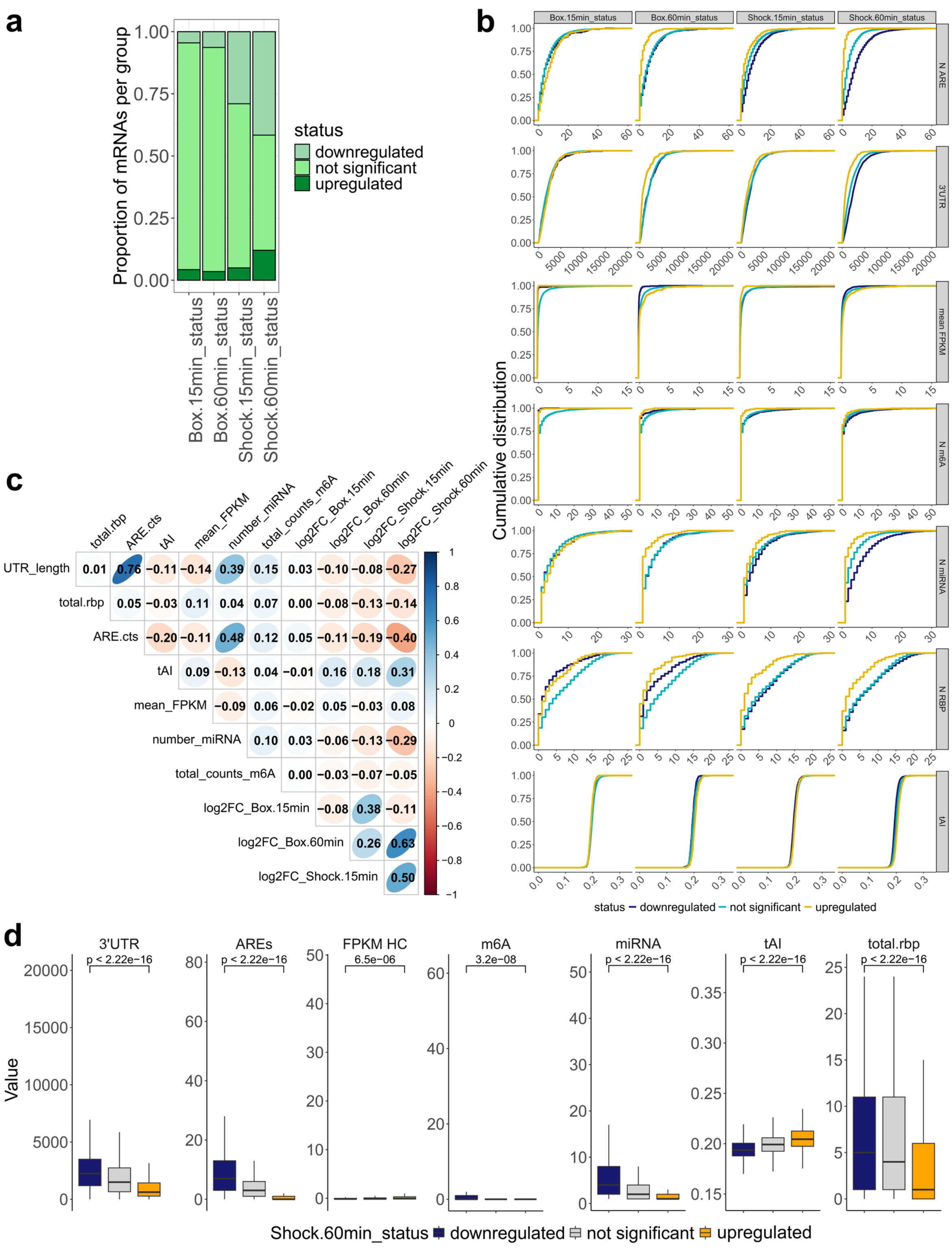


**Ext. Fig 9. Characterization of the input dataset for the *Pvalb*⁺ ML model.** (a) Stacked bar plot illustrating the proportion of genes defined as *downregulated* (top fraction), *not significant* (middle fraction) or *upregulated* (bottom fraction) per group modeled. (b) Cumulative distribution function of each mRNA element, per group modeled. Blue = *downregulated*; Light blue = *not significant*; Orange = *upregulated*. (c) Correlogram illustrating the linear relationship between fold change values of each dataset (normalized to *home cage* controls) and mRNA elements. Numbers inside each square represent the R coefficient. Blue = positive correlation; Red = negative correlation. (d) Box plots directly comparing the prevalence of each mRNA element in *downregulated* versus *upregulated* genes. One-Way ANOVA corrected with Tukey's *post-* [illegible]

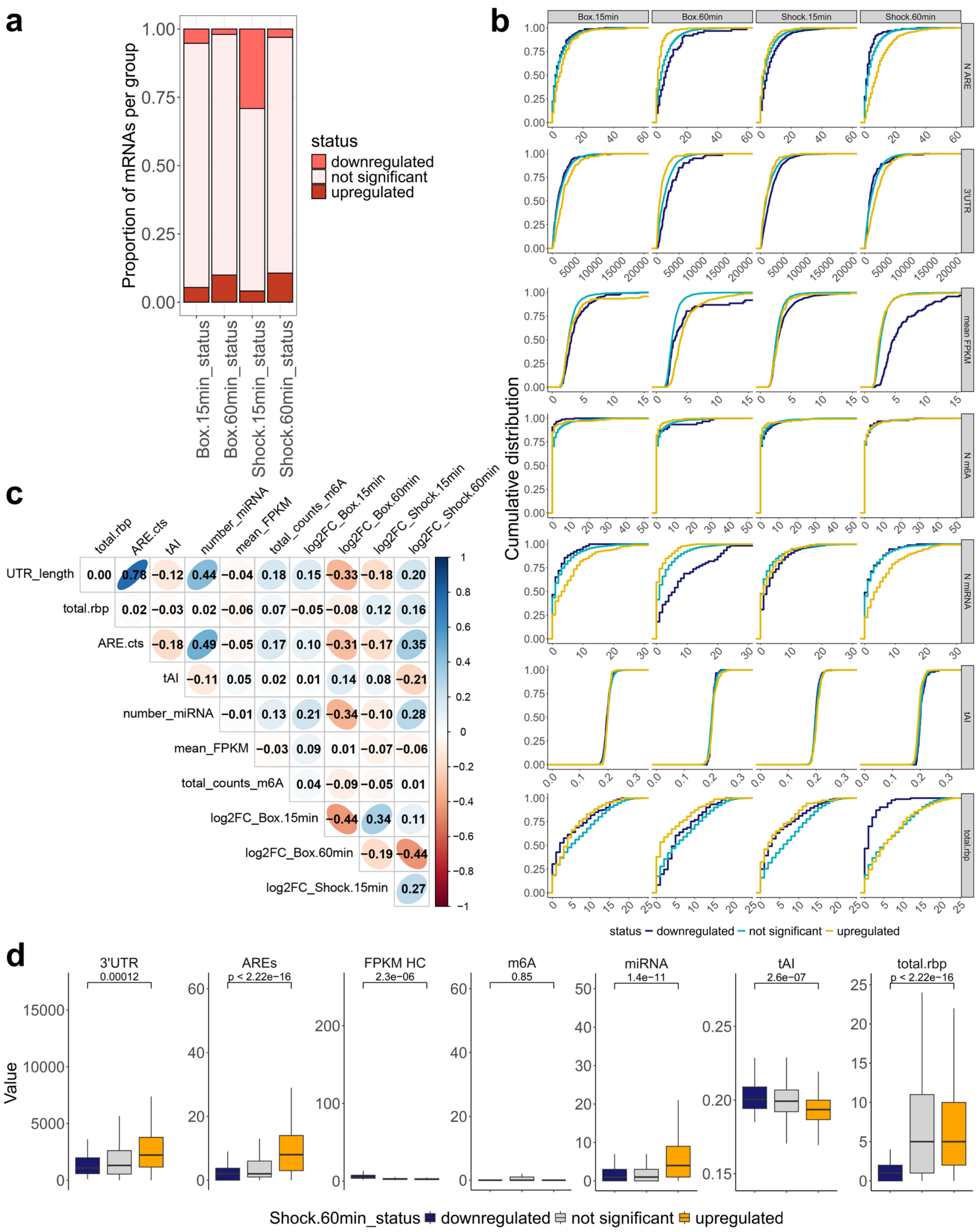


**Ext. Fig 10. Characterization of the input dataset for the *Sst*⁺ ML model.** (a) Stacked bar plot illustrating the proportion of genes defined as *downregulated* (top fraction), *not significant* (middle fraction) or *upregulated* (bottom fraction) per group modeled. (b) Cumulative distribution function of each mRNA element, per group modeled. Blue = *downregulated*; Light blue = *not significant*; Orange = *upregulated*. (c) Correlogram illustrating the linear relationship between fold change values of each dataset (normalized to *home cage* controls) and mRNA elements. Numbers inside each square represent the R coefficient. Blue = positive correlation; Red = negative correlation. (d) Box plots directly comparing the prevalence of each mRNA element in *downregulated* versus *upregulated* genes. One-Way ANOVA corrected with Tukey's *post-hoc* test.

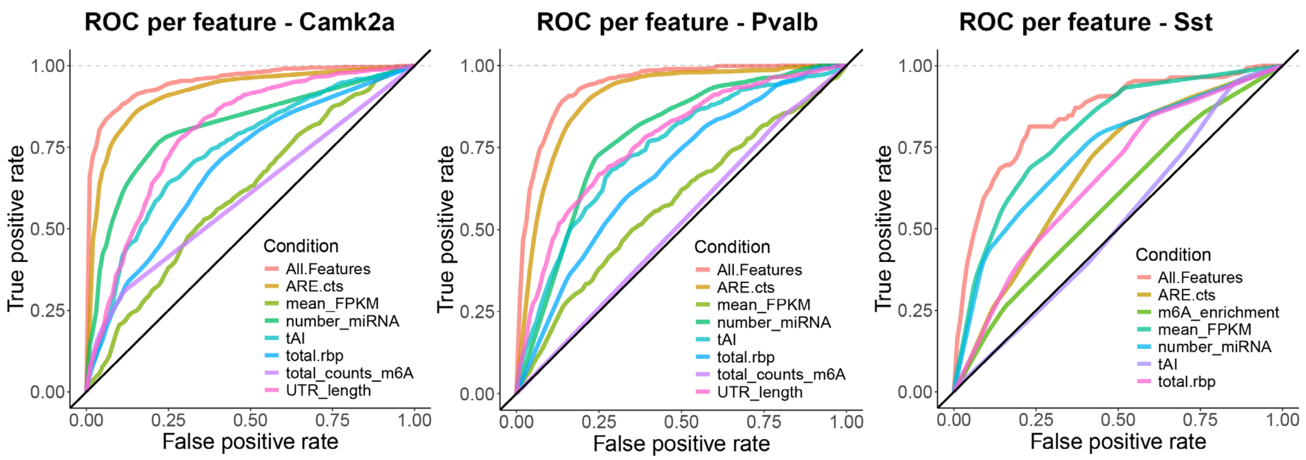


**Ext. Fig 11. Receiving operating characteristic curves for ML models ran with each individual mRNA feature.** From left to right: ROC for *Camk2a*⁺ datasets; ROC for *Pvalb*⁺ datasets; ROC for *Sst*⁺ datasets. Results were obtained running the individual mRNA features only in the 60' shock datasets, as this one demonstrated the highest accuracy across cell types.

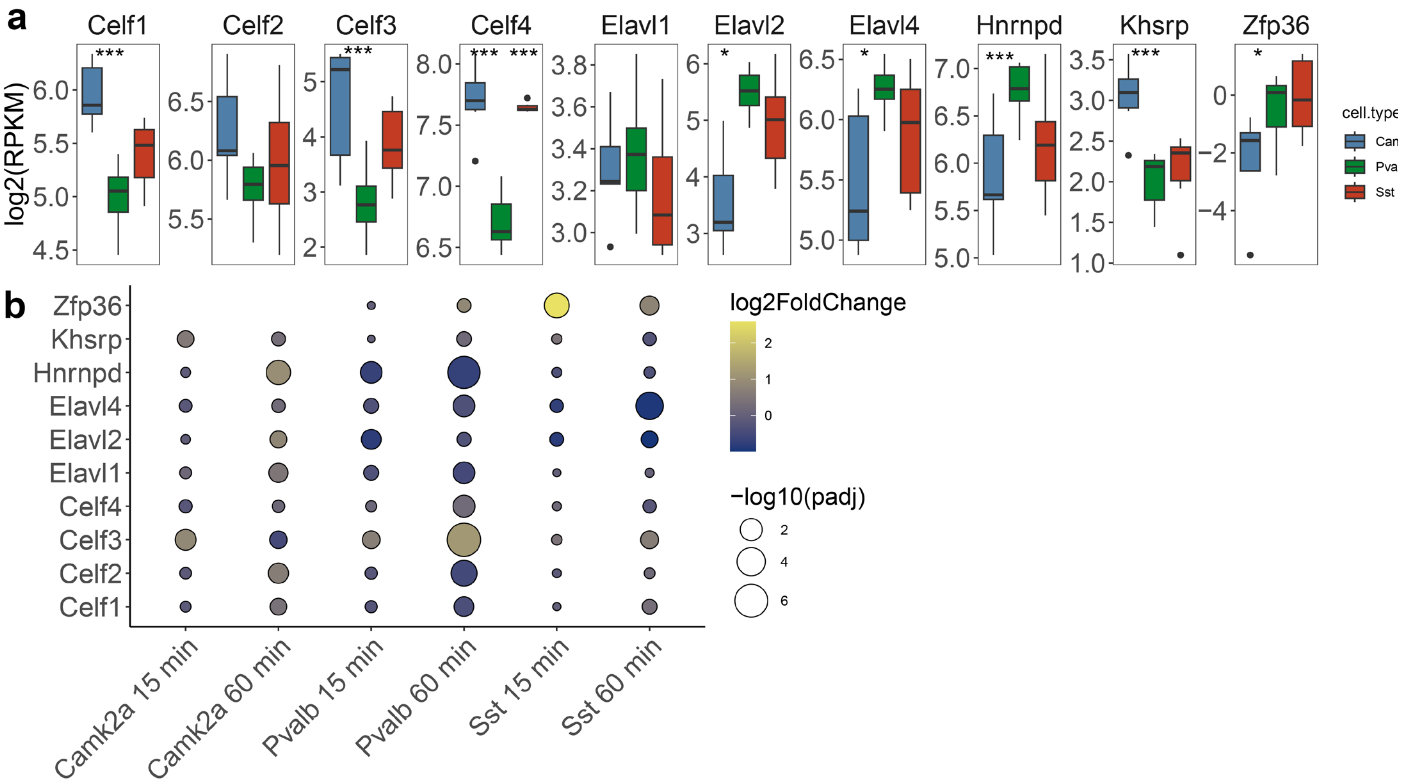


**Ext. Fig 12. Characterization of ARE-targeting RBP expression across different neuron types. (a)** Box plots comparing normalized expression levels (RPKM) of ARE-targeting RBPs between *Camk2a*⁺ (blue), *Pvalb*⁺ (green) and *Sst*⁺ (red) cells. Midline in boxplot represents the median, top and bottom edges represent 75 and 25% quantile ranges, respectively; bars are standard deviation. One-Way ANOVA with Tukey *post-hoc* correction. * = $p < 0.05$; *** = $p < 0.0001$. **(b)** Dot plot showing the differential expression of ARE-targeting RBPs associated with long-term memory consolidation. Color is mapped to log2FC of expression between experimental condition (*Box* or *Shock*) versus baseline control (*Home cage*). Size of circles represent -log10 of *p adj*. Absence of Zfp36 in *Camk2a*⁺ cells is due to lack of significant expression in this cell type (as illustrated in panel (a)).

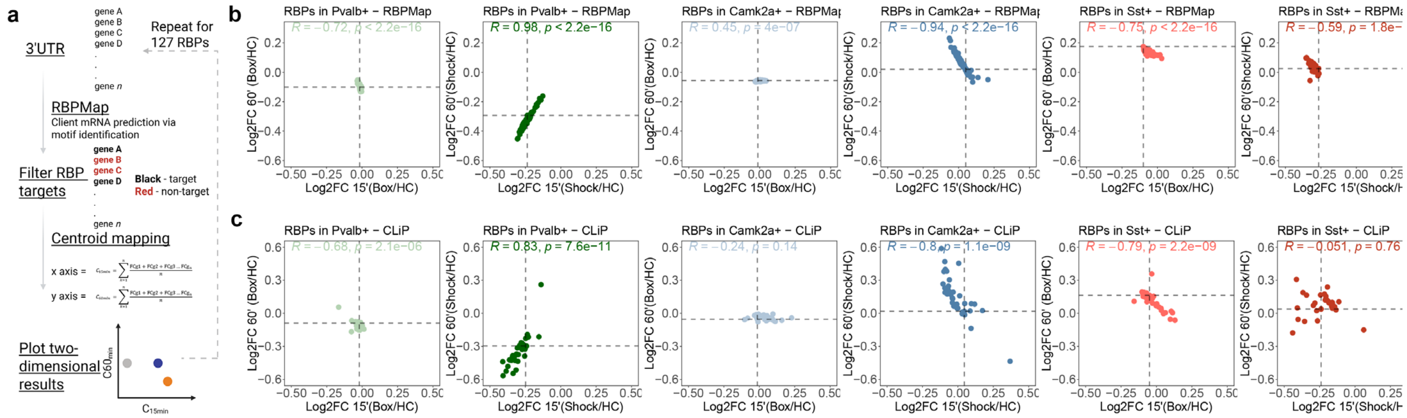

**Ext. Fig 13. Validation of the RBPMap-predicted RBP-RNA interactions database. (a)** Schematics illustrating the process of generating summarizing centroids for each RBP studied. **(b)** Dot plots comparing the time points of 15' (x axis) and 60' (y axis) per condition, per cell type analyzed. Green shades = *Pvalb*$^{+}$ cells datasets; Blue shades = *Camk2a*$^{+}$ cells datasets; Red shades = *Sst*$^{+}$ cells datasets. First, third and fifth columns are representations of the *box* condition, normalized to *home cage*. Second, fourth and sixth columns represent the *shock* condition, normalized to *home cage*. **(c)** results obtained from biological evidence collected using CLiP/metabolic labeling datasets. Represented on top of each plot is the *R* coefficient of a linear regression analysis using the centroid data, comparing the relationship between the two time points analyzed.
